# Cyber-yeast: Automatic synchronisation of the cell cycle in budding yeast through closed-loop feedback control

**DOI:** 10.1101/2020.11.25.398768

**Authors:** Giansimone Perrino, Sara Napolitano, Francesca Galdi, Antonella La Regina, Davide Fiore, Teresa Giuliano, Mario di Bernardo, Diego di Bernardo

**Affiliations:** Telethon Institute of Genetics and Medicine (TIGEM), Pozzuoli, Italy; Department of Chemical, Materials and Industrial Production Engineering, University of Naples Federico II, Naples, Italy; Department of Mathematics and Applications “R. Caccioppoli”, University of Naples Federico II, Naples, Italy; Department of Electrical Engineering and Information Technology, University of Naples Federico II, Naples, Italy; Department of Engineering Mathematics, University of Bristol, Bristol, United Kingdom

## Abstract

The cell cycle is the process by which eukaryotic cells replicate. Yeast cells cycle asynchronously with each cell in the population budding at a different time. Although there are several experimental approaches to “synchronise” cells, these work only in the short-term. Here, we built a cyber-genetic system to achieve long-term synchronisation of the cell population, by interfacing genetically modified yeast cells with a computer by means of microfluidics to dynamically change medium, and a microscope to estimate cell cycle phases of individual cells. The computer implements a “controller” algorithm to decide when, and for how long, to change the growth medium to synchronise the cell-cycle across the population. Our work builds upon solid theoretical foundations provided by Control Engineering. In addition to providing a new avenue for yeast cell cycle synchronisation, our work shows that computers can automatically steer complex biological processes towards desired behaviours similarly to what is currently done with robots and autonomous vehicles.

## INTRODUCTION

The cell cycle is the essential process by which eukaryotic cells replicate. It consists of a sequential series of events that are tightly controlled by an evolutionary conserved regulatory network. The cell cycle acts as a sort of global oscillator regulating cell growth and division^1^. In the budding yeast *S. cerevisiae*, the cell divides asymmetrically with a larger mother budding a smaller daughter cell as outlined in Fig. 1a^2,3^. The cell cycle can be divided into four phases: the growth phase (G1), with a considerable increase in volume, which is followed by the DNA synthesis (S) phase, during which DNA is replicated; afterwards the cell enters a second growth phase (G2), with the appearance of a bud that will grow into the daughter cell, and ends with the mitotic phase (M), when chromosomes become separated and cell division occurs, giving rise to the daughter cell.

**Figure 1.**
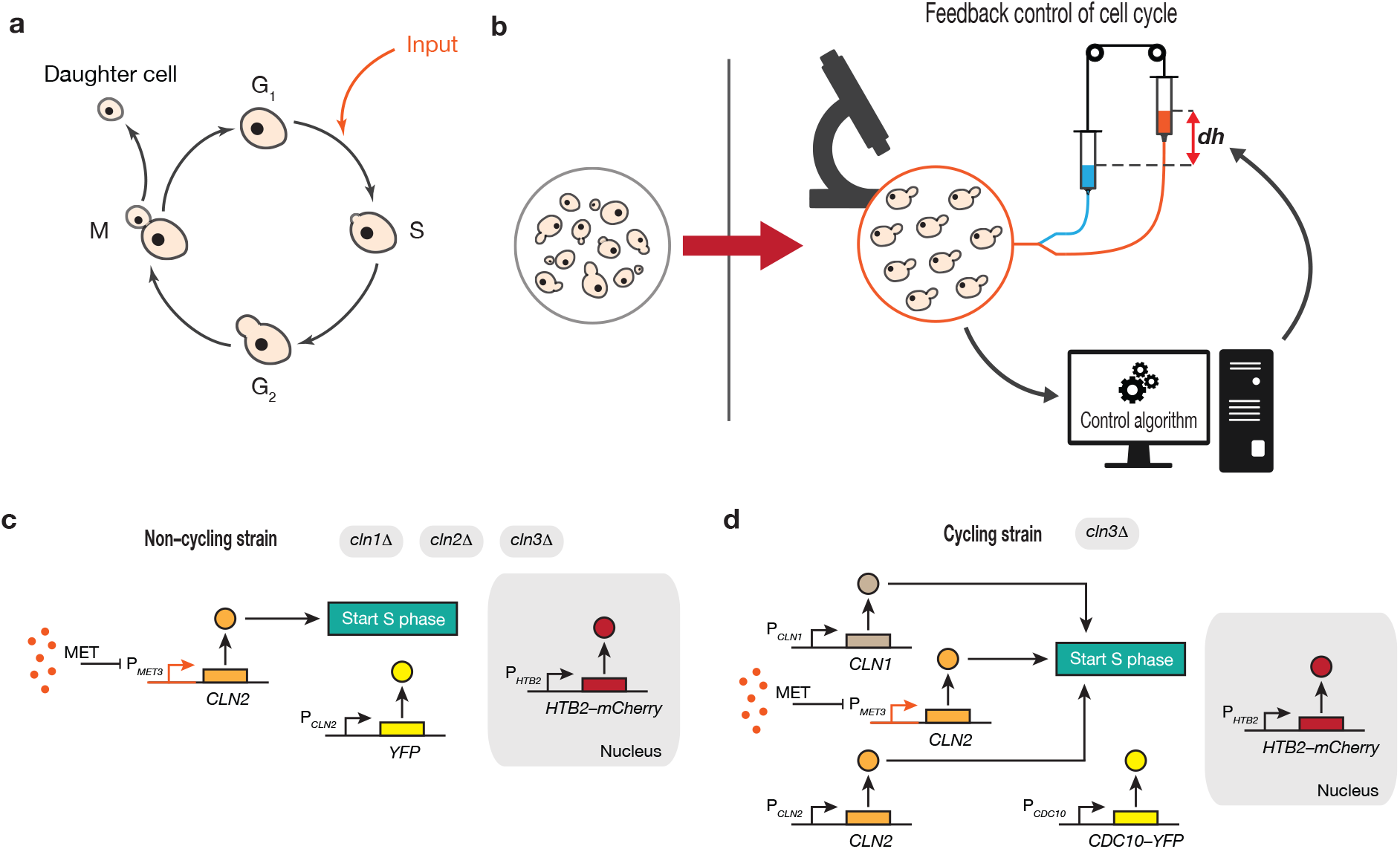
Synchronisation of cell cycle in yeast *S. cerevisiae* through automatic feedback control. **a**, Schematic representation of the cell-cycle in yeast *Saccharomyces cerevisiae*. Yeast strains were engineered to initiate the cell cycle upon methionine depletion from the growth medium (input). **b**, Yeast cells do not cycle synchronously in a population. Schematics of the computer-controlled microfluidics platform to automatically synchronise the cell-cycle across a population of yeast cells. **c**, Non-cycling yeast strain. Cells are deleted for genes encoding for the G_1_ cyclins Cln1-3 while an exogeneous G_1_ cyclin gene *CLN2* is placed under the control of the methionine-repressible promoter P_*MET3*_. Cells can cycle only in the absence of methionine. A yellow fluorescent protein (YFP) is expressed under the control of the endogenous promoter P_*CLN2*_. A red fluorescent nuclear reporter, consisting of a fusion protein between the endogenous histone H2B protein and the mCherry (Htb2-mCherry) is also present in this strain. **d**, Cycling yeast strain. Cells are deleted only for the G_1_ cyclin Cln3, hence cells continuously cycle. An exogeneous copy of the G_1_ cyclin gene *CLN2* is placed under the control of the methionine-repressible promoter P_*MET3*_. A red nuclear fluorescence reporter (Htb2-mCherry) and a yellow fluorescence reporter, consisting of a fusion protein between endogenous mitotic septin Cdc10 and YFP (Cdc10-YFP), are also present in this strain.

During the G_1_ phase, the cell can decide whether to commit to the cell cycle and enter the S phase, or if conditions are not favourable, to arrest the cell cycle. In yeast, this START checkpoint is found in the late G_1_ phase. Activation of the cyclin-dependent kinase (CDK) Cdc28 by any of the cyclins Cln1, Cln2 and Cln3 is necessary to overcome the START checkpoint^4–6^ and to irreversibly enter the cell cycle.

Yeast cells cycle asynchronously, meaning that each cell in the population buds at a different time (Fig. 1b). Such desynchronised behaviour increases cellular heterogeneity and it may be advantageous in unicellular organisms for the survival of the population in unfavourable conditions^7,8^. There are cases, however, where a synchronised population is necessary. For example, the yeast *S. cerevisiae* is the model organism of choice to study the mechanisms underlying eukaryotic cell cycle regulation. This, however, requires a synchronised population of cells to enable robust measurements of transcriptional, proteomic, metabolomic and signalling readouts. Although there are a number of experimental approaches to “synchronise” cells^9–12^, these work only in the short-term, as cells are blocked in the same cell-cycle phase by chemical (e.g. small molecules) or environmental (e.g. temperature) means and then released all together. Hence, after a few generations, cells become again desynchronised^13^. Here, we asked whether we could design an approach to automatically keep the cell cycle synchronised across cells in a population over time.

To tackle this problem, we turned to Control Engineering, a well-established discipline to build “controllers” to regulate the behaviour of physical systems reliably and robustly across a range of operating conditions. “Closed-loop” feedback control is the most common strategy used in Control Engineering to implement a controller. Feedback control relies on a sense and react paradigm (Fig. 1b), where the quantity to be regulated (e.g. the cell-cycle phase of each cell) is measured in real-time and then the controller, usually implemented in a computer, adjusts the input accordingly (e.g. duration and timing of the stimulation) to achieve the control objective (e.g. synchronisation of cell cycle across cells). A key theoretical result of control theory is that feedback control endows systems with robustness to perturbations and uncertainties.

Recently, Charvin et al.^14^ were able to synchronise yeast cells growing in a microfluidics device by inducing periodic expression of the G_1_ cyclin *CLN2* using a methionine-repressible promoter^15^. Among their other findings, they observed that timing of methionine removal and administration (i.e. period and duration of the stimulation) must be carefully tuned to achieve satisfactory synchronisation. Moreover, changes in growth conditions, such as carbon source or temperature, will desynchronise the population, unless the optimal stimulation timing is properly adjusted.

Motivated by these findings, we developed “cyber-yeast”: a completely automated microfluidics platform implementing a feedback control strategy able to maintain a yeast population synchronised over time despite changes in cellular and environmental conditions, as shown in Fig. 1b.

## RESULTS

### Yeast strains for inducible cell cycle start

We made use of two different yeast strains genetically engineered to start the cell cycle upon removal of methionine from the growth medium and shown in Fig. 1c,d.

The first strain in Fig. 1c, which we referred to as the *non-cycling strain*, was engineered by Rahi et al.^16^ (Methods). In the non-cycling strain, endogenous control of cell cycle initiation is disrupted by deletion of the genes encoding for the G_1_ cyclins Cln1, Cln2, and Cln3^17^, whereas *CLN2* is placed downstream of the methionine-repressible promoter P_*MET3*_ to allow its inducible expression^15^. A yellow fluorescence protein (YFP) is expressed from the endogenous P_*CLN2*_ promoter and thus peaks in the late G_1_ phase (Methods). Finally, a constitutively expressed histone Htb2-mCherry acts as a nuclear fluorescence marker for facilitating image analysis (Methods). The non-cycling strain is blocked in the G_1_ phase when grown in methionine-rich medium.

The second strain, which we call the *cycling strain*, is shown in Fig. 1d and it was derived from the one described by Charvin et al.^14^. In this strain, endogenous control of cell cycle initiation is maintained by preserving the genes encoding the G_1_ cyclins Cln1 and Cln2 and by deleting Cln3^17^. An extra copy of *CLN2* is placed under the control of the methionine-repressible promoter P_*MET3*_. Two fluorescence markers are present in this strain: the mitotic septin Cdc10-YFP, expressed during the S-G_2_-M phases, and the Htb2-mCherry as a constitutive nuclear marker (Methods). The cycling strain can cycle independently of methionine levels in the growth medium, however, the cell cycle can also be initiated on demand by inducing exogeneous *CLN2* expression via methionine removal.

### Automatic feedback control of cell cycle in the non-cycling yeast strain

We first experimentally characterised the non-cycling strain. To this end, we grew cells in a microfluidics platform to dynamically change the micro-environment while measuring in real-time fluorescence reporters in individual cells by microscopy, as shown in Fig. 1b (Methods). In methionine-rich medium cells grow in volume but fail to divide for at least 6 hours (Supplementary Fig. 1a-c,g); indeed, some cells were able to replicate after this time, albeit very slowly, possibly because of accumulation of *CLN2* caused by promoter’s leakiness.

In methionine-depleted medium (Fig. 2a-e), cells are able to cycle as evidenced by the exponential increase in cell number over time (Fig. 2a and Supplementary Fig. 1g) and by the cyclic expression of the YFP reporter in individual cells (Fig. 2c, Supplementary Fig. 1d-f, and Supplementary Movie 1). Interestingly, the cell YFP fluorescence intensity averaged across the population shows a flat profile (Fig. 2b), despite being oscillatory in individual cells. This behaviour stems from cells not cycling in phase, thus causing individual oscillations in fluorescence to cancel out when computing the average intensity. Finally, we estimated the budding index (B.I.) from the YFP fluorescence intensities (Methods). The B.I. represents the percentage of cells in the budded phase (i.e. S-G_2_-M phases) at each time point (Fig. 2d) and it is expected to remain constant over time for an unsynchronised cell population. This experimental characterisation of the non-cycling yeast strain thus confirmed that it behaved as expected.

**Figure 2.**
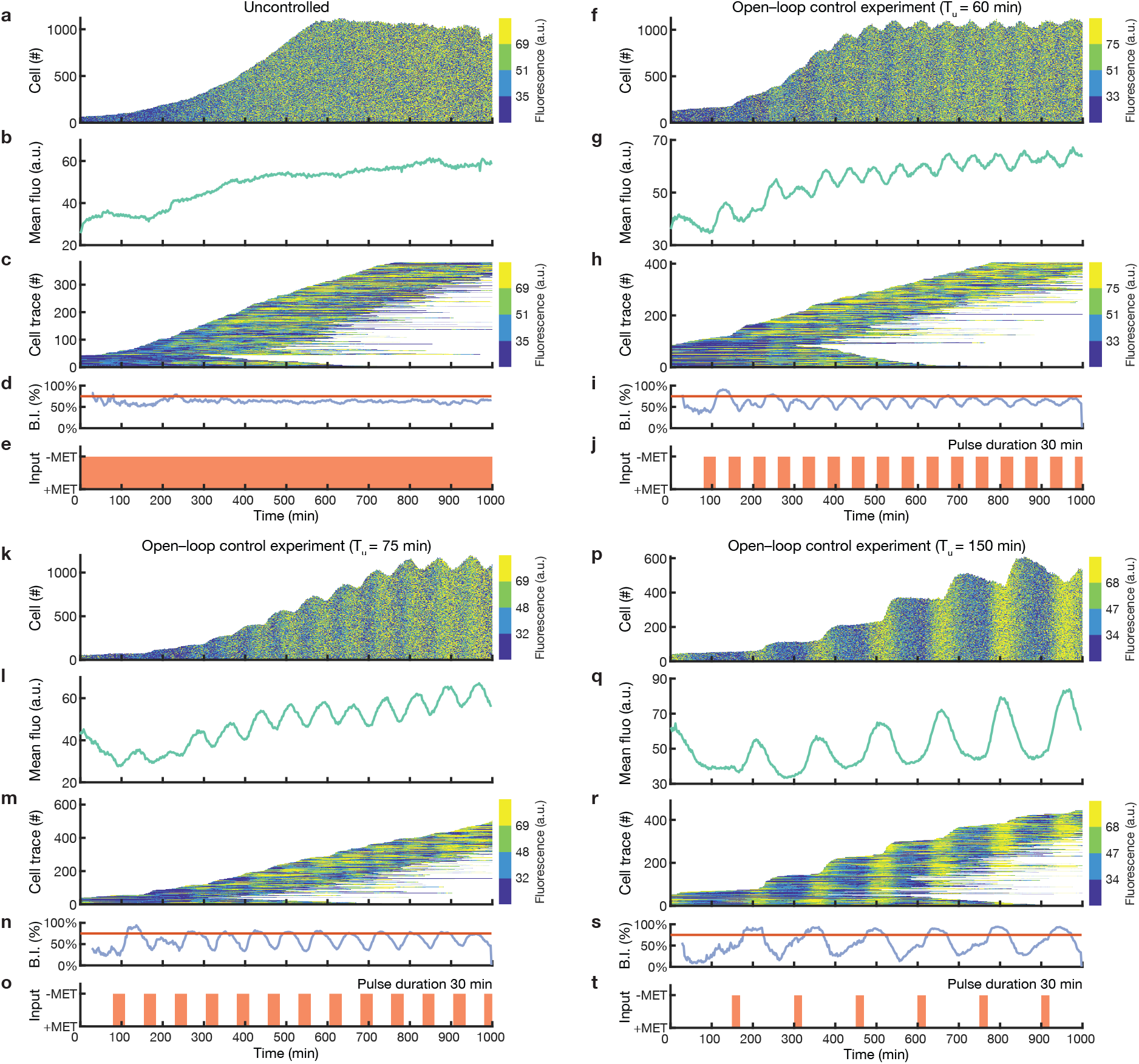
Characterisation and open-loop control of the *non-cycling* yeast strain. Experiments with non-cycling yeast strain cells grown in the automated microfluidics platform in four different conditions: (**a-e**) methionine-depleted medium (-MET); (**f-j**) alternating pulses of methionine-rich (+MET) and methionine-depleted (-MET) medium with a period *T*_*u*_= 60 min; (**k-o**) alternating pulses with a period *T*_*u*_= 75 min; (**p-t**) alternating pulses with a period *T*_*u*_ = 150. Duration of -MET pulse was set to 30 min. **a, f, k, p**, The number of cells and the distribution of YFP fluorescence intensity in the population over time. Fluorescence values are binned into 4 colours, corresponding to the quartiles, for clarity of visualisation. **b, g, l, q**, Average YFP fluorescence intensity in the cell population. **c, h, m, r**, Single-cell fluorescence traces over time. Each horizontal line corresponds to one cell. Each line starts when the cell is first detected and ends when the cell exits the field of view. The number of tracked cells does not correspond to the total number of cells as only cells tracked for longer than 300 min are shown. **d, i, n, s**, Budding index (blue) reporting the percentage of cells in the budding phase (S-G_2_-M) computed from the estimated cell cycle phases. The red line denotes the expected value of the budding index in the case of a totally desynchronised cell population. **e, j, o, t**, Growth medium delivered to the cells as a function of time: +MET methionine-rich medium, -MET: methionine-depleted medium.

Next, we asked whether periodic expression of the G_1_ cyclin *CLN2* in response to periodic pulses of methionine depletion (-MET) would cause individual cells to cycle synchronously. To choose the period (*T*_*u*_) and duration (*D*_−*Met*_) of such pulses, we first derived and analysed a deterministic mathematical model of the cell cycle (Methods). Briefly, the cell cycle was modelled as a phase oscillator, which can be depicted as a clock with a single moving arm whose position indicates the phase of the cell and whose length is proportional to the cell volume (Supplementary Fig. 2a). To simplify the model, we assumed that volume growth in the mother cell occurs only in the G_1_ phase, whereas the bud grows in volume only during the S-G_2_-M phases. In the model, the cell will stop in the G_1_ phase as long as methionine is present in the medium and will jump to the S phase in its absence, but only if its volume is above a critical threshold. The cell cycle duration in the absence of methionine was set to 75 min based on the literature^18^. To simulate a growing population of yeast cells, we generated an agent-based model where each cell is a phase oscillator, and a new agent is added after each cell cycle is completed. We first simulated the mathematical model to predict the behaviour of the cell population in response to external periodic expression of *CLN2* varying the stimulation period *T*_*u*_ and the pulse duration *D*_−*Met*_ (Supplementary Fig. 2b,d). Numerical simulations show that the forcing period *T*_*u*_ is of paramount importance for achieving cell cycle synchronisation across the cell population. Indeed, to fully synchronise the population, the period *T*_*u*_ must be greater than the intrinsic cell cycle period (i.e. *T*_*u*_ ≥ 75 min) (Supplementary Fig. 2b). Moreover, the longer the period (*T*_*u*_), the larger the average cell volume (*V*) (Supplementary Fig. 2d), as cells stay in the G_1_ phase for longer.

Guided by these numerical results, we performed microfluidics-based experiments to periodically induce the expression of *CLN2* with -MET pulses of period *T*_*u*_ varying between 60 min and 150 min, and duration *D*_−*Met*_ = 20 min or 30 min. Experimental results are consistent with numerical simulations, as both synchronisation of the cell cycle across the population and the cells’ average volume increase with the period *T*_*u*_ (Fig. 2f-t, Supplementary Fig. 3 and Supplementary Movies 2 and 3). The fluorescence intensities of cells over time, in Fig. 2f,k and p, show a clear vertical pattern, indicating that cells are mostly in the same cell cycle phase; additionally, the number of cells increases in a step-wise function, rather than exponentially, as most of the cells bud together. Furthermore, the population averaged YFP fluorescence intensity (Fig. 2g,l and q) displays an oscillatory behaviour, as cells become synchronised. A similar result was obtained by computing the budding index (Fig. 2i,n and s) from single-cell traces (Fig. 2h,m and r). Finally, we observed that for a shorter duration of - MET pulses (*D*_−*Met*_ = 20 min and *T*_*u*_ = 75 min), cells’ synchronisation takes considerably longer (700 min) as compared to a longer pulse (*D*_−*Met*_ = 30 min and *T*_*u*_ = 75 min), (Supplementary Fig. 3 and Fig. 2k-o). The average cell radius and the extent of synchronisation were quantified for each experiment and reported in Supplementary Fig. 4 by evaluating both the mean coherence phase *R* and the amplitude of the leading peak in the power spectrum of the average fluorescence signal.

**Figure 3.**
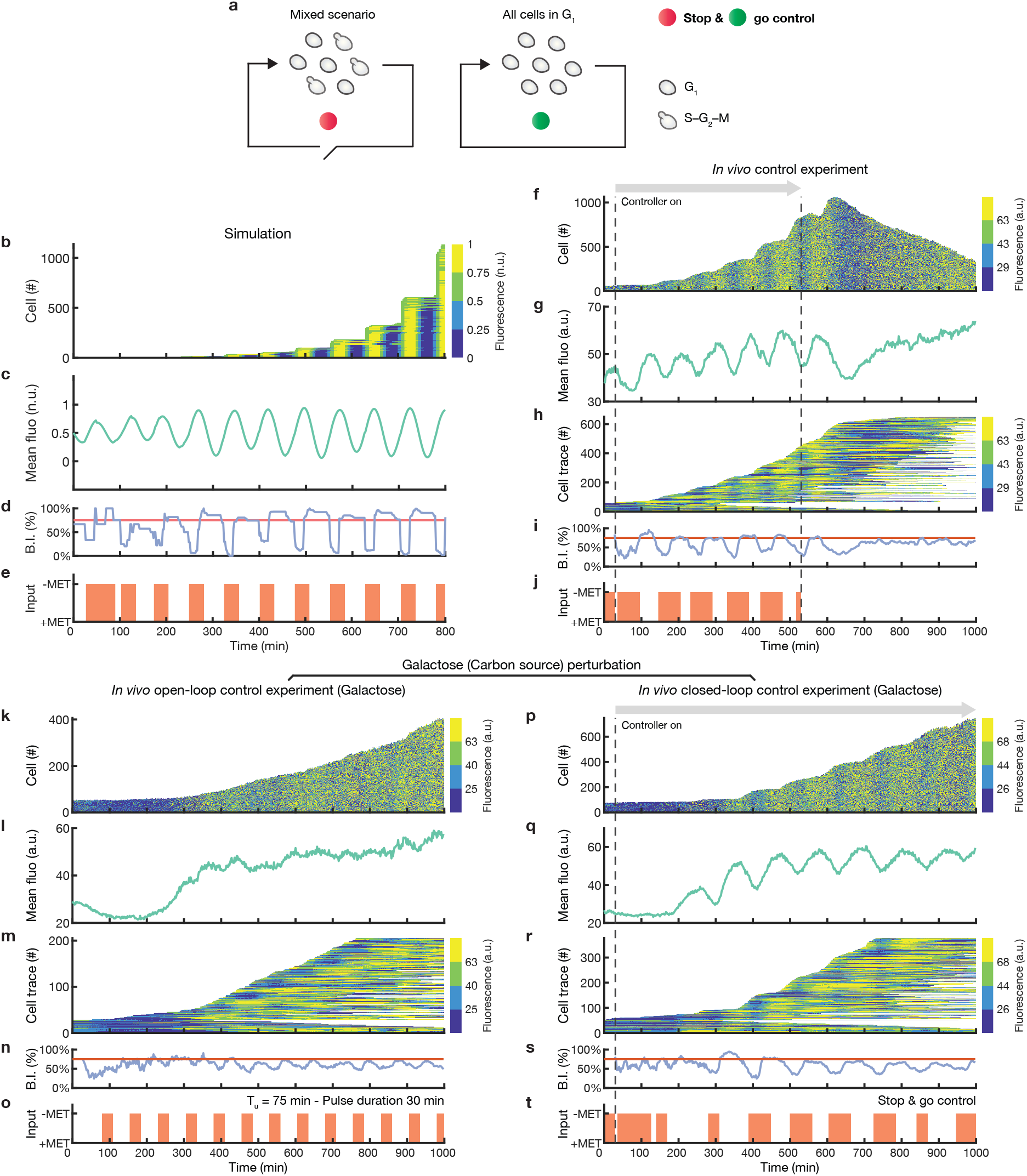
Automatic feedback control enables cell cycle synchronisation in the *non-cycling* yeast strain. **a**, Depiction of the *stop & go* control strategy. The controller waits for cells to stop in the G_1_ phase and then restarts their cell cycle by removing methionine from the medium. **b-e**, Numerical simulation of the stop & go control strategy. **f-j**, Experimental implementation of the stop & go control strategy. An initial calibration phase of 30 min was required to set up the phase estimation algorithm. Dashed lines indicate the start and the end of the control experiment, after which cells are grown in methionine-rich medium. **k-o**, Open-loop experiment subjecting cells to alternating pulses of methionine-rich (+MET) and methionine-depleted (-MET) medium with a period *T*_*u*_ = 75 min using galactose (2% w/v) as carbon source. **p-t**, Experimental implementation of the stop & go control strategy using galactose (2% w/v) as carbon source. **b, f, k, p**, The number of cells and the distribution of YFP fluorescence intensity in the population over time. Fluorescence values are binned into 4 colours, corresponding to the quartiles, for clarity of visualisation. **c, g, l, q**, Average YFP fluorescence intensity in the cell population. **h, m, r**, Single-cell fluorescence traces over time. Each horizontal line corresponds to one cell. Each line starts when the cell is first detected and ends when the cell exits the field of view. The number of tracked cells does not correspond to the total number of cells as only cells tracked for longer than 300 min are shown. **d, i, n, s**, Budding index (blue) reporting the percentage of cells in the budding phase (S-G_2_-M) computed from the estimated cell cycle phases. The red line denotes the expected value of the budding index in the case of a totally desynchronised cell population. **e, j, o, t**, Growth medium delivered to the cells as a function of time: +MET methionine-rich medium, -MET: methionine-depleted medium.

**Figure 4.**
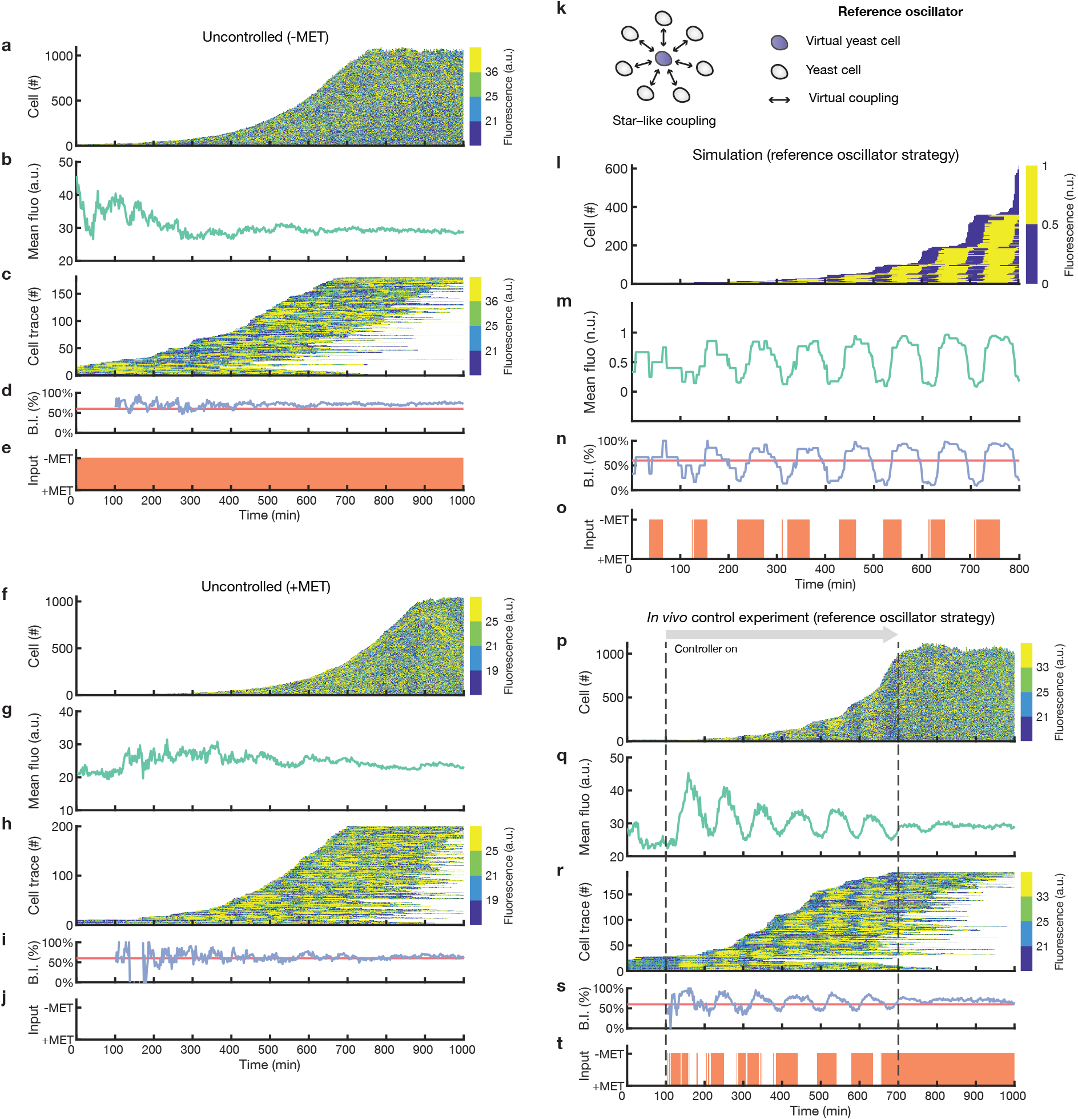
Automatic feedback control enables cell cycle synchronisation of the *cycling* yeast strain. **a-j**, Experimental characterisation of the cycling yeast strain grown in the automated microfluidics platform in (**a-e**) methionine-depleted (-MET) medium or (**f-j**) methionine-rich medium (+MET). **k**, Schematic illustration of the reference oscillator control strategy. Yeast cells are coupled to a *virtual* cell so that each cell cycles in sync with the virtual cell. **l-o**, Numerical simulation of the reference oscillator control strategy. **p-t**, Experimental implementation of the reference oscillator control strategy. An initial calibration phase of 100 min is required to set up the phase estimation algorithm. Dashed lines indicate the start and the end of the control experiment, after which cells are grown in methionine-depleted medium. **a, f, l, p**, The number of cells and the distribution of YFP fluorescence intensity in the population over time. Fluorescence values are binned into 4 colours, corresponding to the quartiles, for clarity of visualisation. **b, g, m, q**, Average YFP fluorescence intensity in the cell population. **c, h, r**, Single-cell fluorescence traces over time. Each horizontal line corresponds to one cell. Each line starts when the cell is first detected and ends when the cell exits the field of view. The number of tracked cells does not correspond to the total number of cells as only cells tracked for longer than 300 min are shown. **d, i, n, s**, Budding index (blue) reporting the percentage of cells in the budding phase (S-G_2_-M) computed from estimated cell cycle phases. Red line denotes the expected value of the budding index in the case of a totally desynchronised cell population. **e, j, o, t**, Growth medium delivered to the cells as a function of time: +MET methionine-rich medium, -MET: methionine-depleted medium.

Notwithstanding the feasibility of synchronising cells using periodic stimulation, this “open-loop” control strategy is highly sensitive to environmental conditions, for example whereas periodic stimulation with period *T*_*u*_ = 75 min and pulse-duration *D*_−*Met*_ = 30 min is able to synchronise the cell cycle when cells are grown in nominal conditions (glucose at 30°C in Fig. 2k-o), this is not the case when growing cells in galactose at 30°C (Fig. 3k-o) or in glucose at 27°C (Supplementary Fig. 5a-e).

To overcome the drawbacks of the open-loop approach, we implemented a “closed-loop” feedback control strategy to automatically synchronise the cell population. To this end, we exploited the characteristics of the non-cycling strain to be blocked in the G_1_ phase in methionine-rich medium (+MET) and to cycle upon its depletion (-MET). We devised an event-triggered “stop & go” control strategy (Methods), whose working principle is illustrated in Fig. 3a: at each sampling time, the controller estimates the cell cycle phase of each cell across the population from fluorescence microscopy images (Methods). If the percentage of cells in the G_1_ phase is higher than a fixed threshold, then the controller delivers an exogenous pulse of -MET, thus enabling the cell cycle to start in all G_1_ phase cells.

To test the feasibility of the stop & go control strategy in achieving the synchronisation task, we first carried out a numerical simulation with a threshold value set to 50% (Fig. 3b-e). Numerical results show that the stop & go controller can in principle synchronise the cell cycle across a population of cells, without any prior knowledge of the cell cycle duration and growth conditions. Figure 3b shows the simulated fluorescence signal in individual cells over time, with clear vertical lines and a step-wise increase in cell number indicating synchronisation; the population average fluorescence signal (Fig. 3c) and the budding index (Fig. 3d) are oscillatory, as expected for synchronised cells. Interestingly, after a transient, the external input became a quasi-periodic signal (Fig. 3e) and entrained the single-cell traces with a 1:1 ratio (Fig. 3b). Finally, we carried out numerical simulations for different values of the threshold used by stop & go controller to decide when to deliver the -MET pulse. As shown in Supplementary Fig. 2c,e, the larger the threshold, the more the cells become synchronised, but at the cost of larger cell volumes.

Encouraged by the numerical results, we experimentally implemented the stop & go feedback control strategy with a threshold set to 50%. Cells were grown overnight in the absence of methionine so that, at the beginning of each experiment, the population was totally desynchronised (Methods). An initial calibration phase of 30 min was required to initialise the phase estimation algorithm (Methods), after that the stop & go controller was activated for 500 min and then deactivated. The closed-loop stop & go control strategy can automatically synchronise the cell population, as shown in Fig. 3f-j, Supplementary Fig. 6a-j and Supplementary Movie 4. Fluorescence intensities including all of the cells, (Fig. 3f; Supplementary Fig. 6a,f) and single-cell fluorescence traces, available only for a subset of cells for which cell tracking was successful, (Fig. 3h; Supplementary Fig. 6c,h) show a clear vertical pattern over time, indicating synchronisation, while the cell number increases in a step-wise fashion, as long as the controller is active. In addition, the average fluorescence signal and the budding index show an oscillatory behaviour (Fig. 3g,i; Supplementary Fig. 6b,d,g,i). As observed in simulations, after a transient, the control input became a quasi-periodic signal (Fig. 3j; Supplementary Fig. 6e,j). Quantification of average cell radius and the extent of synchronisation are reported in Supplementary Fig. 4.

To check the robustness of the stop-and-go feedback controller and to compare it to the open-loop control strategy, we performed additional perturbation experiments by either changing the growth temperature or the carbon source. When the cell growth temperature was set to 27°C (Supplementary Fig. 5), the closed-loop controller was still able to synchronise the cell population (Supplementary Fig. 5f-j) with the same performance as in the unperturbed case (Supplementary Fig. 4), unlike the open-loop periodic stimulation strategy (Supplementary Fig. 5a-e and Supplementary Fig. 4). Next, we performed closed-loop control with cells growing in galactose, which is known to slow down the cell cycle^19^. As shown in Fig. 3k-t, and quantified in Supplementary Fig. 4, the closed-loop feedback control achieved synchronisation without any drop in performance, whereas open-loop periodic stimulation failed to synchronise the population.

Taken together, our results demonstrate that closed-loop feedback control can automatically synchronise yeast cells over time, even in the face of new and unexpected environmental perturbations.

### Automatic feedback control of cell cycle in a cycling yeast strain

Next, we investigated cell cycle synchronisation in the cycling yeast strain in Fig. 1d. The main difference with respect to the non-cycling strain is that cells can cycle also in the presence of methionine. Nevertheless, when in the G_1_ phase, cells can be forced to start the cell cycle by removing methionine from the medium. Synchronisation of the cell population is much more challenging than in the non-cycling strain, as the cells continuously cycle and can never be blocked. As an analogy, imagine a group of basketball players, each constantly moving while bouncing a ball, the challenge is to synchronise the players so that all the balls touch the ground at the same time. If the players are allowed to hold the ball still (non-cycling strain) than it suffices to ask the players to hold the ball and release it at the same time (stop-and-go strategy), however if the players are not allowed to hold the ball (cycling strain) then the solution is much more difficult.

We first performed a series of time-lapse microfluidics experiments to assess cell cycle dynamics both in the absence (Fig. 4a-e) and in the presence of methionine (Fig. 4f-j and Supplementary Movie 5). As expected in both conditions, the number of cells increases exponentially over time and cells are totally desynchronised (Supplementary Fig. 4). Charvin et al. have previously demonstrated that, in this strain, periodic expression of cyclin *CLN2* by alternating pulses of methionine depletion can induce partial synchronisation of cell cycle across the population^14^. They also showed that synchronisation is very sensitive to the period of stimulation^14^.

Here, we asked whether a closed-loop feedback control strategy can improve synchronisation performance and robustness in this strain. The stop-and-go strategy is obviously not applicable, thus at first we implemented a Model Predictive Control (MPC) strategy, which has been successfully applied to biological control problems^20–24^ (Methods). To this end, we modified the agent-based mathematical model derived for the non–cycling strain to adapt it to the cycling strain (Methods). The working principle of the MPC strategy is illustrated in Supplementary Fig. 7a. Briefly, the controller uses the mathematical model of the cell cycle in each cell to predict the future behaviour of the population of cycling cells in order to find the best sequence of methionine-depletion pulses that has to be applied to maximise cell cycle synchronisation. To test the feasibility of the MPC strategy in achieving the synchronisation task, we first carried out a numerical simulation using the deterministic mathematical model (Supplementary Fig. 7b-e). Simulation results show that the MPC controller can partially synchronise the cell population.

We next tested the MPC strategy experimentally and assessed the synchronisation performance, as reported in Supplementary Fig. 7f-t. During microfluidics experiments, the computer took measurements at 2 min intervals and calculated the optimal sequence of stimulation to deliver at 100 min intervals. In each experiment, cells were grown overnight in methionine-rich medium and were totally desynchronised (Methods). An initial calibration phase of 100 min is required to set up the phase estimation algorithm (Methods). We first tested the MPC strategy by performing a closed-loop control experiment for 900 min (Supplementary Fig. 7f-j); partial synchronisation of the cell cycle was achieved, although with a poor performance as evidenced by the low-amplitude oscillations in the average YFP fluorescence intensity and in the budding index in Supplementary Fig. 7g,i and by the quantitative analysis (Supplementary Fig. 4). We confirmed these results by performing two additional experiments (Supplementary Fig. 7k-t) in which the controller was turned off after 600 min. These results demonstrated the feasibility to synchronise the cell cycle using the MPC strategy, but they also highlighted its main drawbacks, i.e. limited performance, high computational costs and the need of a model, which makes the MPC less robust to biological uncertainties.

We thus asked whether a simpler model-independent feedback control strategy could be found with increased performance and reduced computational cost. To this end, we implemented the *reference oscillator* feedback control strategy, which was adapted from the one proposed by Bai and Wen to synchronise a homogenous population of nonlinear phase oscillators using a common input^25^. At the core of this strategy lies a continuously cycling computer-generated *in silico* virtual yeast cell coupled to the real cells via the microfluidics device according to a star-like topology (Fig. 4k). In this context, the virtual cell behaves as a reference oscillator for all the other cells. Using the basketball players analogy, this is like projecting a virtual player on the wall and asking the real players to bounce the balls in synchrony with the virtual player. Mathematical details of this strategy are described in the Methods section. Numerical simulation of the reference oscillator strategy in Fig. 4l-o shows effective synchronisation of the cell cycle population with superior performance with respect to the MPC strategy (Supplementary Fig. 7b-e) despite reduced computational costs and dispensing with modelling of individual cells.

Experimental results for the reference oscillator feedback control strategy are shown in Fig. 4p-t, Supplementary Fig. 8a-j and Supplementary Movie 6. The experimental conditions were the same as for the MPC experiments. The controller was able to quickly synchronise the cell population in agreement with numerical simulations, whereas once the controller was turned off, the cells quickly desynchronised. Moreover, quantitative comparison of the performance of the MPC and reference oscillator strategy in Supplementary Fig. 4 confirms the superiority of the latter strategy.

These results demonstrate that automatic feedback control can synchronise the cell cycle in a population of cycling yeast cells.

## DISCUSSION

A defining feature of yeast is its ability to bud daughter cells by following a precise sequence of events. Each cell within a population, however, buds at a different time (unsynchronised). Biologists and biotechnologists have long searched for methods to synchronise cells, but only short-term synchronisation has been achieved so far^13^. Here, we built a completely automatic system able to achieve long-term synchronisation of the cell population, by interfacing genetically modified yeast cells with a computer by means of microfluidics to dynamically change medium, and a microscope to estimate cell cycle phases of individual cells. The computer implements a “controller” algorithm to decide when, and for how long, to change the growth medium to synchronise the cell-cycle across the population. Our work builds upon solid theoretical foundations provided by Control Engineering. This blending of disciplines is at the core of the new field of Cybergenetics that aims at augmenting biological systems with controllers to regulate their behaviour for biotechnological and biomedical applications^26^.

In addition to providing a new avenue for yeast cell cycle synchronisation, our work shows that classical control engineering, originally devised for steering the behaviour of physical systems, can be successfully applied also to complex biological processes such as the cell cycle. The robustness intrinsic to closed-loop feedback control enables synchronisation to occur in a variety of environmental conditions (i.e. carbon source, temperature), unlike open-loop approaches such as periodic administration of methionine, which are very sensitive to environmental fluctuations.

Our work also lays the basis for engineering the first self-synchronising yeast strain: this could be achieved by implementing the stop & go controller genetically within each cell, rather than using a computer, by employing synthetic biology approaches^27^.

## METHODS

### Yeast strains and growth conditions

All *Saccharomyces cerevisiae* strains used in this study are listed in Supplementary Table 1. Strains are congenic with W303 strain. The SJR14a4d strain^16^ (a kind gift from S. J. Rahi) is the non-cycling strain. The yDdB028 strain was derived from the GC84-35B strain^14^ (a kind gift from G. Charvin) and represents the cycling strain.

The yDdB028 strain was constructed using standard procedures. The *HTB2-mCherry* cassette was amplified via PCR on genomic DNA extracted from the SJR14a4d strain and cloned into plasmid pRS41N-GAP-CYC (a derivative of the nourseothricin-selective pRS41N plasmid^28^ containing the CYC terminator, a kind gift from C. Wilson). The plasmid sequence was checked by Sanger sequencing. The *HTB2-mCherry* cassette was amplified from plasmid pRS41N-GAP-HTB2-mCherry-CYC and inserted into the yeast HTB2 locus via homologous recombination^29^ such that expression was driven by the endogenous promoter. Correct integration was verified by PCR on extracted genomic DNA. All plasmids used in this study are listed in Supplementary Table 2.

Unless otherwise specified, both strains were grown at 30°C in either synthetic complete medium, composed of yeast nitrogen base (0.67% w/v) with all amino acids; or synthetic complete drop-out medium, composed of yeast nitrogen base (0.67% w/v) with all amino acids except methionine; both supplemented with glucose (2% w/v) as carbon source. For the carbon source perturbation experiments, synthetic complete media were supplemented with galactose (2% w/v).

### Microfluidics

All microfluidics experiments were performed with the MFD0005a device^30^. This device contains a chamber in which the cells are trapped. The height of the chamber (3.5 μm) allows the yeast cells to grow only in a monolayer, simplifying the image analysis. Microfluidics devices were fabricated with a replica moulding technique as previously described^21^. The fluid that reaches the chamber of the microfluidics device is a mixture of the growth media coming from the two inlet ports. The blending of the growth media depends on the relative pressure between the two fluids at the inlet ports. In order to change the relative pressure, we devised an automated actuation system that varies the relative height of the two syringes filled with the +MET and -MET media. The actuation system relies on two custom vertically mounted linear actuators, that can move independently. Each of the two linear actuators comprises one stepper motor connected with a syringe through a timing belt and two pulleys. A custom MATLAB script pilots the stepper motors to automatically move the syringes and thus direct the fluid from the one or the other inlet to the cell chamber.

### Microscopy image acquisition and processing

Phase contrast and epifluorescence images were acquired at 2 min intervals at 40 × magnification (CFI Plan Fluor DLL 40 × dry objective, NA 0.75; Nikon Instruments) using a Nikon Eclipse Ti-E inverted microscope (Nikon Instruments) coupled with an EMCCD cooled camera (iXon Ultra 897; Andor Technology). The microscope stage was surrounded by an opaque cage incubator (Okolab) able to maintain the temperature at either 30°C (nominal condition) or 27°C (perturbed condition). Time-lapse experiments were conducted with the Perfect Focus System (Nikon Instruments) enabled. Appropriate filter cubes were used to acquire the yellow (YFP HYQ and FITC for the cycling and the non-cycling strains, respectively; Nikon Instruments) and the red (TRITC HYQ; Nikon Instruments) fluorescence channels. Time-lapse image acquisition was controlled by the NIS-Elements Advanced Research software (Nikon Instruments).

Raw phase contrast and epifluorescence images were processed using custom scripts implemented in the MATLAB environment available here (https://github.com/dibbelab/Cycloop). Briefly, the images in the red channel were used to identify single cell nuclei visible thanks to the nuclear fluorescence marker (Htb2-mCherry). Next, the nuclei centroids were used as seeds for a Voronoi tessellation to generate a single-cell region mask to crop the phase contrast images around each cell. The resulting phase contrast cropped images were used to generate a binary mask defining the region of a single cell. We applied the function *regionprops* to all the single-cell mask to quantify the radius of each cell. Fluorescence intensities were then quantified by processing the yellow fluorescence images with the binary single-cell masks described above, using the function *regionprops*. Specifically, for the non-cycling strain the fluorescence is quantified as the average fluorescence intensity in the region selected by the binary mask; while for the cycling strain the fluorescence is quantified as the maximum fluorescence intensity detected in the region selected by the binary mask. Fluorescence intensities are measured in arbitrary units. Single-cell traces were tracked in real-time using a custom tracking algorithm previously described^21^. To improve the single-cell trace datasets, we devised an offline implementation of the tracking module that performs also a reverse tracking of the cell population, which means that the tracking module was run a second time starting however from the last time frame towards the first frame. Thus, we obtained a reverse single-cell traces’ dataset. Then, the algorithm combined the forward and the reverse datasets to improve the single-cell trace dataset.

### Microfluidics experiments

The microfluidics experimental platform was initialised as previously described^22^. For microfluidics experiments performed in nominal conditions, cells from a frozen glycerol stock (−80°C) were resuspended in 10 mL of either methionine-free (non-cycling strain) or methionine-supplemented (cycling strain) growth medium, grown overnight in a shaking incubator at 220 r.p.m. and 30°C, and then injected in the microfluidics device as previously described^22^. Unless otherwise specified, cells were left to settle in the chamber for 15 min fed with either methionine-depleted (non-cycling strain) or methionine-rich (cycling strain) growth medium. After that, the operator run the image acquisition and the custom MATLAB software. At the beginning of the experiment, a *region of interest* (ROI) was selected on the first acquired phase contrast image. Specifically, the ROI defines the area containing the *S. cerevisiae* cells that have to be segmented and tracked, and whose fluorescence signals have to be quantified. For the carbon source perturbation experiments, cells were treated as in the nominal conditions except that galactose was the only carbon source added to the growth media. For the temperature perturbation experiments, cells were grown as in the nominal conditions except that the temperature was maintained at 27°C in lieu of 30°C.

### Modelling

We constructed a deterministic agent-based mathematical model to quantitatively describe the collective dynamical behaviour of cell cycle in both strains (cycling and non-cycling). Our agent-based model is based on previously published models^27,31,32^, where each agent represents a cell in the population. The model of a single agent is based on a set of two state-dependent ordinary differential equations (ODEs), which track the evolution of cell cycle phase *ϑ* and cell volume *V* in each cell:

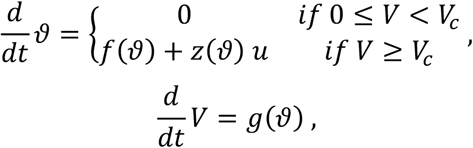

where *ϑ* ∈ 𝕊^1^ is the 2π-periodic cell cycle phase on the unit circle, *V* ∈ ℝ_+_ is the cell volume, *V*_c_ ∈ ℝ_+_ is the critical volume, and *u* ∈ {0,1} is the external trigger input. The critical volume defines the minimum volume required to start the cell cycle, and it is also used to discern between mother (*V* ≥ *V*_*c*_) and daughter (0 ≤ *V* < *V*_*c*_) cells. The external binary input represents the methionine-rich (*u* = 0) or methionine-depleted (*u* = 1) growth medium. The phase-dependent switching function *g* : 𝕊^1^ → ℝ_+_ defines the cell volume growth rate during the G_1_ phase:

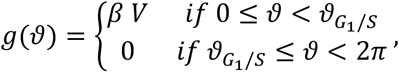

where *β* ∈ ℝ_+_ is the volume growth rate, and 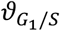 is the cell cycle phase value at the G_1_ to S transition. We assumed that cell’s volume grows exponentially only during the G_1_ phase, thus neglecting the mass generated during the other phases (i.e. S-G_2_-M), most of which is transferred to the growing bud^14^. The phase-dependent switching function *f* : 𝕊^1^ → ℝ_+_ models the phase oscillator dynamics. The function *f* changes according to the specific strain. For the non-cycling strain, the function *f* := *f*_*nc*_ is state-dependent:

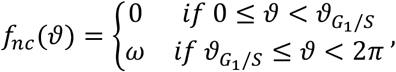

where 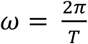 is the angular velocity depending on the cell cycle period *T*. For the cycling strain, the function *f* := *f*_c_ becomes state-independent:

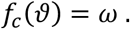

Finally, the phase response curve *z* : 𝕊^1^ → ℝ_+_ models the linear response of the cell cycle phase *ϑ* to the input *u*:

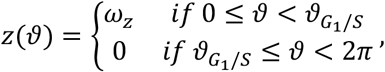

where *ω*_*z*_ ∈ ℝ_+_ is the angular velocity added to the cell cycle phase dynamics when the cell is fed with methionine-free medium.

Our agent-based model also considers cell division events. Indeed, when a cell passes through the M to G_1_ transition 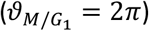 then a new cell (i.e. a new agent) is added to the model. The initial condition of the daughter cell depends on the state of the mother cell. Specifically, the initial phase is set to *ϑ*_1_ = 0, while the initial volume is set to 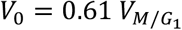, where 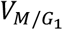 is the volume of the mother cell at the division time (i.e. at the M/G_1_ transition). Therefore, the number of cells in our agent-based model is an increasing value.

The parameter values used in the agent model of the non-cycling strain are: *V*_*c*_ = 1 (critical volume), *β* = 0.0083 *min*^−1^ (volume growth rate; see ref.^14^), *T* = 75 *min* (nominal cell cycle period; see ref.^18^),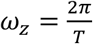. (angular velocity in methionine-free medium), and 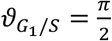. (phase value at G_1_/S transition; see ref.^33^). The parameter values used in the agent model of the cycling strain are: *V*_*c*_ = 1 (critical volume), *β* = 0.0083 *min*^−1^ (volume growth rate; see ref.^14^), *T* = 105 *min* (nominal cell cycle period; see Phase estimation and budding index section), 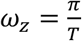 (additional angular velocity in methionine-free medium), and 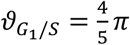 (phase value at G_1_/S transition; see Phase estimation and budding index section). All simulations were performed using an initial population of *N*_0_ = 3 homogeneous cells. Initial phases were uniformly spaced in the interval [0, 2*π*[. Initial volumes were set equal to the critical volume *V*_*c*_. The agent-based system was solved using the MATLAB *ode15s* solver. All plots were generated in MATLAB.

### Phase estimation and budding index

For the non-cycling strain, the cell cycle phase *ϑ* ∈ [0, 2*π*] was estimated by comparing the single-cell *CLN2-YFP* trace with a periodic reference signal. The periodic reference signal *CLN*2_*ref*_ was constructed according the dynamical expression of the essential G_1_ cycling gene *CLN2* in the non-cycling strain:

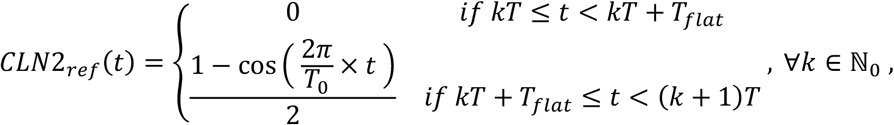

where *T* = *T*_*flat*_ + *T*_0_ is the period of the reference signal, *T*_*flat*_ is a fake time interval in which the cell is assumed to be halted in the G_1_ phase, and *T*_0_ is the nominal cell cycle period in the non-cycling strain. The first part of the periodic reference signal describes the situation in which the cell has a flat fluorescence signal, that is when it is halted in the G_1_ phase, and thus the duration *T*_*flat*_ must be set equal to the length of the compared fluorescence signal. Instead, the second part models the oscillatory *CLN2-YFP* expression in a cell that is cycling, i.e. when it is fed with methionine-free medium. At each sampling time *t*, the measured fluorescence signal was cross-correlated with the periodic reference signal evaluating the Pearson’s correlation coefficient *r*. Specifically, the last part (duration equals to *T*_*flat*_) of the measured fluorescence signal 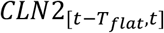 was cross-correlated with the periodic reference signal using a shifting time window in the interval [*τ* − *T*_*flat*_),*τ*], where *τ* ∈ [*T*_*flat*_, *T*]. The time point *τ* at which reached the maximum value was used as a time-reference to estimate the cell cycle phase 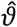 using the linear relationship:

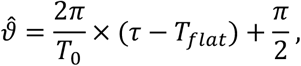

where 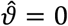 means that the cell is at the M/G_1_ transition. The Pearson’s correlation coefficient was computed using the MATLAB function *corr*. The nominal cell-cycle period *T*_0_ was set to 75 min, i.e. the duration of the cell cycle period in the non-cycling strain^18^; while the time period *T*_*flat*_ was set to 30 min, according the length of the measured fluorescence signal used to evaluate the cross-correlation. The length of the G_1_ phase was set to the 25% of the nominal cell cycle period^33^.

For the cycling strain, the cell cycle phase *ϑ* ∈ [0, 2*π*] was estimated from the single-cell *CDC10-YFP* trace using a custom procedure. The single-cell fluorescence signal was firstly binarized according the dynamical expression of the septine protein Cdc10. Indeed, the septine protein Cdc10 contributes to form the ring at the interface between the mother and the daughter cell (i.e. the bud)^34^. Thus, the fluorescence signal is linked to the bud formation and so is detected only during the budded phase (i.e. S-G_2_-M phases). In the ideal case, the fluorescence signal is binary (on/off). To binarize the single-cell fluorescence trace at the sampling time *t*, the raw Cdc10-YFP signal was coarsely smoothed with a moving average filter with window size of 5 samples and then filtered again with a finer 3 samples window filter. Then, the filtered fluorescence signal was transformed in a binary signal using the threshold value *l*(*t*):

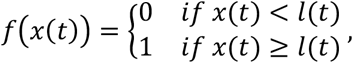

where *x* is the filtered fluorescence value, and *l* is the threshold value; both evaluated at time *t*. *f*(*x*) = 0 means that the cell is in the off state, i.e. the unbudded phase; while *f*(*x*) = 1 means that the cell is in the on state, i.e. the budded phase. The threshold value *l* at time *t* was computed from the raw fluorescence data extracted from the final 100 min of the acquired single-cell fluorescence signals. Specifically, the raw fluorescence values for all the cells were aggregated together in a single dataset, and then a mixture of two normal distributions was fitted on the histogram of this dataset. The threshold level was chosen as the intersection point of the two normal distributions. Next, the single-cell binary signal was checked to detect the last binary transition. This can be an on to off (*on* → *off*) transition, corresponding to the start of the unbudded phase; or an off to on (*τ* → *on*) transition, corresponding to the start of the budded phase. Assuming that the last transition occurred at time *t* – *T*_*t*_, the estimated cell cycle phase 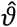 was computed as:

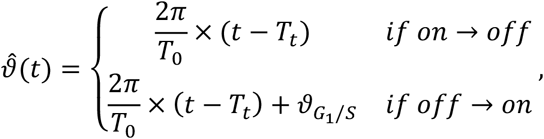

where *T*_0_ is the nominal cell cycle period in the cycling strain, *T*_*t*_ is the time interval from the last binary transition to the current time, and 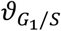 is the phase value at which occurs the G_1_/S transition. If a cell did not show a binary transition during the observation period, then the cell cycle phase could not be estimated. To further improve the quality of results, we set a minimum duration of 14 min between two successive binary transitions. Part of this procedure was used to estimate the nominal cell cycle period *T*_0_ and the phase value 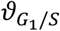 from the raw fluorescence traces of an uncontrolled +MET experiment (Supplementary Data 1). Specifically, the single-cell raw fluorescence signals were binarized using a threshold value *l*. Then, the nominal duration of the unbudded phase *T*_*G*1_ = 42 min was computed as the median of the intervals measured in all binary traces between an on to off (*pn* → *off*) and an off to on (*off* → *on*) transition. Similarly, the nominal duration of the budded phase 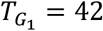 min was computed as the median of the intervals measured in all binary traces between an off to on (*off* → *on*) and an on to off (*on* → *off*) transition. Finally, the numerical values of *T*_1_ and *ϑ*_*G*_1__/*_S_* were obtained as:

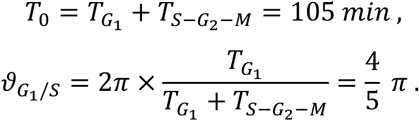

The budding index (B.I.) was computed as the percentage of budded cells in the population:

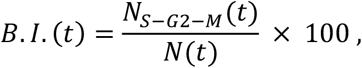

where *N*(*t*) is the total number of cells at time *t*, and *N*_*S* − *G*2 − *M*_(*t*) is the number of budded cells (i.e. cells in the S-G_2_-M phases) at time *t*.

### Quantification of synchronization

To quantify the degree of cell cycle synchronisation across the cell population, we computed two synchronisation indices: a) the mean coherence phase *R* of the Kuramoto order parameter^35^ and b) the amplitude of the leading peak in the power spectrum of the average YFP fluorescence signal.

The mean phase coherence *R*∈ [0,1] is defined as the magnitude of the Kuramoto order parameter^35^:

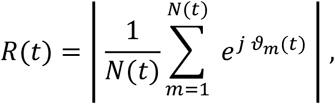

where *N*(*t*) is the time-varying number of cells, while *ϑ*_*m*_(*t*) is the phase of the *m*-th cell, both evaluated at time *t*. When *R* is equal to 1, all cells are synchronised. Conversely, when *R* is equal to 0, cells are totally desynchronised.

The amplitude of the leading peak was computed from the power spectral density (PSD) of the normalised YFP fluorescence signal. Specifically, the mean of the average YFP signal was subtracted from the signal itself. Then the MATLAB function *pmcov* of the *Signal Processing Toolbox*, was used to computer a parametric estimation of the PSD. The leading peak of the PSD was found with the MATLAB function *max*. The leading peak in the PSD is associated to a period. We decided to discard a peak value associated to a period longer than the duration of the YFP signal. In this case, we classified that peak as not detectable (N.D.).

### Control algorithms

All control algorithms were devised to change the control input (i.e., the growth medium) once a new system output (i.e., the estimated cell cycle phases) was available. Moreover, the control input was held constant between two consecutive phase measurement times (zero-order hold method^36^). All control algorithms were implemented through custom scripts in the MATLAB environment.

#### Stop-and-go control

The stop-and-go control algorithm is an event-triggered feedback control strategy that was devised to synchronising the cell cycle across a population of non-cycling cells^31^. At each sampling time *t*_*k*_, the stop-and-go algorithm computes the percentage of cells in the G_1_ phase by means of the estimated cell cycle phases (see Phase estimation and budding index section). If the percentage of cells in the G_1_ phase is higher than a fixed threshold *v*_%_, then the algorithm delivers a - MET pulse of duration *D*_−*Met*_ to the cells, otherwise cells are kept in +MET medium. Controller’s parameters used in our implementation were: *v*_%_ = 50% (threshold value) and *D*_−*Met*_ = 30 *min* (duration of -MET pulse). Note that *v*_%_ = 100% in the ideal implementation illustrated in Fig. 3a.

#### Model predictive control (MPC)

The MPC algorithm is an optimization-based feedback control strategy that was devised to synchronise the cell cycle across a population of cycling cells^32^. Specifically, the MPC algorithm was designed to solve an open-loop optimal control problem repeatedly over a receding horizon using the deterministic agent-based mathematical model described in the Modelling section^37^. This means that at each iteration of the algorithm, the solution to the optimal open-loop control problem gives an optimal control input (i.e., the sequence of -MET pulses) that maximises the cell cycle synchronisation over a finite prediction horizon *T*_*p*_. The optimal control input is applied only over a finite control horizon *T*_*c*_ ≤ *T*_*p*_. In our implementation, both the prediction horizon *T*_*p*_ and the control horizon *T*_*c*_ were chosen equal to 100 min. To reduce the computational complexity of the optimal control problem, we made a series of assumptions on both the agent-based mathematical model and the control input: (i) we neglected both the cell growth and the cell division in the model of each cell agent; (ii) we assumed that the -MET pulse can reset the phase *ϑ* to the value 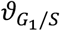 (i.e., the G_1_ to S phase transition) when the cell is in G_1_ phase. Thus, the mathematical model of a single agent was reduced to one ODE, which describes the evolution of the cell cycle phase *ϑ* in each cell:

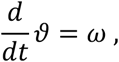

where 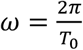 is the angular velocity depending on the cell cycle period *T*. Let *u* ∈ {0, 1} be the external ideal trigger input to the agent model, where *u* = 1 corresponds to methionine-depleted medium and *u* = 0 to methionine-supplemented medium. The effect of the control input on the cell cycle phase evolution was modelled with the following phase reset rule:

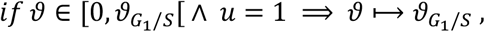

where Λ is the notation adopted for the *and* operator. Furthermore, the control *u* ∈ 𝒰 was assumed to be a finite sequence of ideal triggers. Defining *P* ∈ ℕ as the maximum number of triggers that may be applied in a finite prediction horizon [*t*_*k*_, *t*_*k*_ + *T*_*p*_[, then the time interval between two consecutive triggers was 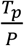. The feasible set *u* was defined as the set of all admissible control sequences and was thus composed by 2^*p*^ possible combinations of triggers. In our implementation, we considered *P* = 10, that means 1024 possible combinations of triggers in the prediction horizon. To reduce again the computational complexity, we made a series of assumptions on the admissible sequences of triggers. Specifically, we imposed that the -MET pulse duration *D*_− *Met*_ should be constrained into the interval 20 ≤ *D*_−*Met*_ ≤ 60 *min*. To avoid loss of synchronization due to a prolonged absence of stimulation, we also imposed that the maximum time distance between two consecutive -MET pulses should be equal to 60 min. At each iteration of the MPC algorithm, we considered a fixed number of homogeneous agents *N* equals to the number of cell cycle phases estimated by the phase estimation algorithm in that sampling time (see Phase estimation and budding index section). The estimated cell cycle phases were also used to set the initial conditions of the deterministic agent-based model. In the model of each agent, the model parameter *T*_0_ was set equal to the nominal value reported in the Modelling section. The optimal control problem was solved by maximising a synchronisation index *J* over the prediction horizon *T*_*p*_. Specifically, the synchronisation index *J* ∈ [0, 1] quantified the cell cycle synchronisation over time in each simulation:

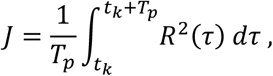

where *R* is the mean phase coherence of the Kuramoto order parameter computed with the simulated cell cycle phases (see Quantification of synchronisation section). Optimisation was achieved by exploring all the possible admissible combinations of -MET pulse sequences.

#### Reference oscillator

The reference oscillator strategy is a state feedback control strategy that was devised to synchronise the cell cycle across a population of cells. We adapted the control strategy proposed by Bai and Wen^25^ to design a state feedback control law able to steer the cell cycle phases to converge towards a reference phase. Here, the reference phase evolves over time according to the dynamics of a virtual phase oscillator that interacts with all the cells in the population through a virtual star-like coupling. We constructed the virtual reference oscillator through an ODE model that describes the evolution of the reference oscillator phase *ϑ*_*r*_ on the unit circle 𝕊^1^ over time:

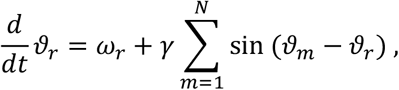

where *ω*_*r*_ ∈ ℝ_+_ is the natural frequency of the reference oscillator, *γ* ∈ ℝ_+_ is the coupling strength, *N* ∈ ℕ is the number of cells, and *ϑ*_*m*_ ∈ 𝕊^1^ is the cell cycle phase of the *m*-th cell in the population. Note that the number of cells changes over time at each sampling time *t*_*k*_ (see Phase estimation and budding index section). Moreover, the cell cycle phases were held constant between two consecutive phase measurement times (zero-order hold method^36^). At each sampling time *t*_*k*_, the algorithm first measures the cell cycle phases through the phase estimation algorithm (see Phase estimation and budding index section). Let *N*_C_ be the number ofcells whose phases have been measured at the sampling time *t*_*k*_. Then, the algorithm computes the control action *u*(*t*_*k*_) to apply to the cell population in the interval [*t*_*k*_, *t*_*k*+1_[through the state feedback control law:

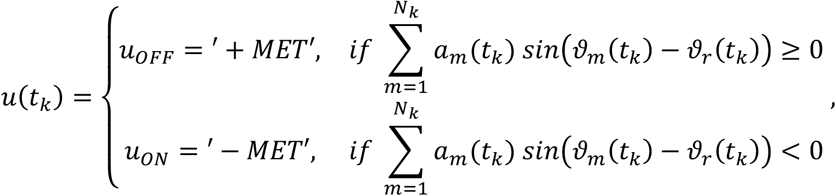

where *ϑ*_*r*_(*t*_*k*_) is the phase of the reference oscillator at the sampling time *t*_*k*_, *ϑ*_*m*_(*t*_*k*_) is the cell cycle phase of the *m*-th cell measured at the sampling time *t*_*k*_; and *a*_*m*_(*t*_*k*_) ∈ {0,1} is a time-dependent coefficient associated to the *m*-th cell denoting if that cell is in the G_1_ phase at the sampling time *t*_*k*_, that is:

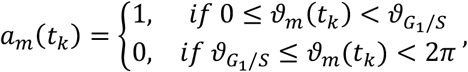

where 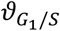 is the phase at the G_1_ to S transition according the phase estimation algorithm (see Phase estimation and budding index). Finally, the algorithm integrates the ODE model of the reference oscillator to obtain the value of the reference phase *ϑ*_*r*_ at the next sampling time *t*_*k*+1_. To integrate the reference oscillator ODE, we used the MATLAB numerical solver *ode45*. Controller’s parameters used in our implementation were: *ω*_*r*_ equals to the nominal natural velocity of the cell cycle in the cycling strain (see Modelling section), and γ = 1 (coupling strength). The initial condition of the reference oscillator phase was *ϑ*_*r,0*_ = 0. For the complete details of the reference oscillator strategy, including the derivation of the state feedback control law, refer to the Supplementary Notes.

## Data availability

All relevant data generated in this study are deposited at Zenodo (https://doi.org/10.5281/zenodo.4045689). Strains and plasmids used in this study are available from the corresponding author upon reasonable request.

## Code availability

The source code that supports the findings of this study is available at GitHub (https://github.com/dibbelab/Cycloop).

## ACKNOWLEDGMENTS

The authors would like to thank Sahand Jamal Rahi (EPFL, Switzerland) for the SJR14a4d *S. cerevisiae* strain (i.e. the non-cycling strain), Gilles Charvin (IGBMC, France) for the GC84-35B *S. cerevisiae* strain (i.e. background of the cycling strain), Cathal Wilson (TIGEM, Italy) for the insightful help with strain construction and for the plasmid pRS41N-GAP-CYC, and Jeff Hasty (UCSD, USA) for the master mould of the microfluidic device. This work was supported by the Fondazione Telethon grant TGM16SB1 and by the COSY-BIO (Control Engineering of Biological Systems for Reliable Synthetic Biology Applications) project, which has received funding from the European Union’s Horizon 2020 research and innovation programme under grant agreement 766840.

## AUTHOR CONTRIBUTIONS

G.P., S.N. and D.d.B. designed the research; G.P. developed and supervised all the controllers’ designs and modelling; S.N. performed and supervised the experiments and developed image analysis and phase estimation algorithms; F.G. helped to design the MPC controller and the phase estimation algorithms; A.L.R. helped to perform experiments, image analysis and modelling; D.F. helped to develop the Stop & Go controller and the cell cycle model; T.G. engineered the cycling yeast strain; M.d.B. helped in controllers’ design and modelling; G.P., S.N. and D.d.B. analysed the data; D.d.B. proposed the concept and supervised the project; G.P., S.N. and D.d.B. wrote the manuscript.

## REFERENCES

1. Li, F., Long, T., Lu, Y., Ouyang, Q. & Tang, C. The yeast cell-cycle network is robustly designed. Proc. Natl. Acad. Sci. U. S. A. 101, 4781 LP–4786 (2004).

2. Morgan, D. O. The cell cycle: principles of control. (New science press, 2007).

3. Hartwell, L. H. & Unger, M. W. Unequal division in Saccharomyces cerevisiae and its implications for the control of cell division. J. Cell Biol. 75, 422–435 (1977).

4. Dirick, L., Böhm, T. & Nasmyth, K. Roles and regulation of Cln-Cdc28 kinases at the start of the cell cycle of Saccharomyces cerevisiae. EMBO J. 14, 4803–4813 (1995).

5. Tyers, M., Tokiwa, G. & Futcher, B. Comparison of the Saccharomyces cerevisiae G1 cyclins: Cln3 may be an upstream activator of Cln1, Cln2 and other cyclins. EMBO J. 12, 1955–1968 (1993).

6. Cross, F. R. Starting the cell cycle: what’s the point? Curr. Opin. Cell Biol. 7, 790–797 (1995).

7. Acar, M., Mettetal, J. T. & van Oudenaarden, A. Stochastic switching as a survival strategy in fluctuating environments. Nat. Genet. 40, 471–475 (2008).

8. Beaumont, H. J. E., Gallie, J., Kost, C., Ferguson, G. C. & Rainey, P. B. Experimental evolution of bet hedging. Nature 462, 90–93 (2009).

9. Juanes, M. A. Methods of Synchronization of Yeast Cells for the Analysis of Cell Cycle Progression. Methods Mol. Biol. 1505, 19–34 (2017).

10. Hur, J. Y., Park, M. C., Suh, K.-Y. & Park, S.-H. Synchronization of cell cycle of Saccharomyces cerevisiae by using a cell chip platform. Mol. Cells 32, 483–488 (2011).

11. Davis, P. K., Ho, A. & Dowdy, S. F. Biological Methods for Cell-Cycle Synchronization of Mammalian Cells. Biotechniques 30, 1322–1331 (2001).

12. Williamson, D. H. & Scopes, A. W. A Rapid Method for Synchronizing Division in the Yeast, Saccharomyces Cerevisiae. Nature 193, 256–257 (1962).

13. Talia, S. Di, Skotheim, J. M., Bean, J. M., Siggia, E. D. & Cross, F. R. The effects of molecular noise and size control on variability in the budding yeast cell cycle. Nature 448, 947–951 (2007).

14. Charvin, G., Cross, F. R. & Siggia, E. D. Forced periodic expression of G1 cyclins phase-locks the budding yeast cell cycle. Proc. Natl. Acad. Sci. 106, 6632--6637 (2009).

15. Amon, A., Irniger, S. & Nasmyth, K. Closing the cell cycle circle in yeast: G2 cyclin proteolysis initiated at mitosis persists until the activation of G1 cyclins in the next cycle. Cell 77, 1037–1050 (1994).

16. Rahi, S. J., Pecani, K., Ondracka, A., Oikonomou, C. & Cross, F. R. The CDK-APC/C Oscillator Predominantly Entrains Periodic Cell-Cycle Transcription. Cell 165, 475–487 (2016).

17. Skotheim, J. M., Di Talia, S., Siggia, E. D. & Cross, F. R. Positive feedback of G1 cyclins ensures coherent cell cycle entry. Nature 454, 291–296 (2008).

18. Rahi, S. J. et al.. Oscillatory stimuli differentiate adapting circuit topologies. Nat. Methods 14, 1010–1016 (2017).

19. Nguyen-Huu, T. D. et al.. Timing and Variability of Galactose Metabolic Gene Activation Depend on the Rate of Environmental Change. PLOS Comput. Biol. 11, e1004399 (2015).

20. Postiglione, L. et al.. Regulation of Gene Expression and Signaling Pathway Activity in Mammalian Cells by Automated Microfluidics Feedback Control. ACS Synth. Biol. 7, 2558–2565 (2018).

21. Perrino, G., Wilson, C., Santorelli, M. & di Bernardo, D. Quantitative Characterization of α-Synuclein Aggregation in Living Cells through Automated Microfluidics Feedback Control. Cell Rep. 27, 916-927.e5 (2019).

22. Fiore, G., Perrino, G., di Bernardo, M. & di Bernardo, D. In Vivo Real-Time Control of Gene Expression: A Comparative Analysis of Feedback Control Strategies in Yeast. ACS Synth. Biol. 5, 154–162 (2016).

23. Uhlendorf, J. et al.. Long-term model predictive control of gene expression at the population and single-cell levels. Proc. Natl. Acad. Sci. 109, 14271 LP–14276 (2012).

24. Milias-Argeitis, A. et al.. In silico feedback for in vivo regulation of a gene expression circuit. Nat. Biotechnol. 29, 1114–1116 (2011).

25. Bai, H. & Wen, J. T. Asymptotic Synchronization of Phase Oscillators With a Single Input. IEEE Trans. Automat. Contr. 64, 1611–1618 (2019).

26. Khammash, M., Bernardo, M. D. & Bernardo, D. Di. Cybergenetics: Theory and Methods for Genetic Control System. In 2019 IEEE 58th Conference on Decision and Control (CDC) 916–926 (2019).

27. Perrino, G. & di Bernardo, D. Synchronisation of yeast cell cycle through quorum sensing coupling. bioRxiv (2020) doi:10.1101/2020.04.05.026179.

28. Taxis, C. & Knop, M. System of centromeric, episomal, and integrative vectors based on drug resistance markers for Saccharomyces cerevisiae. Biotechniques 40, 73–78 (2006).

29. Gardner, J. M. & Jaspersen, S. L. Manipulating the yeast genome: deletion, mutation, and tagging by PCR. Methods Mol. Biol. 1205, 45–78 (2014).

30. Ferry, M. S., Razinkov, I. A. & Hasty, J. Microfluidics for synthetic biology: from design to execution. Methods Enzymol. 497, 295–372 (2011).

31. Perrino, G., Fiore, D., Napolitano, S., Bernardo, M. di & Bernardo, D. di. Towards feedback control of the cell-cycle across a population of yeast cells. In 2019 18th European Control Conference (ECC) 2644–2650 (2019).

32. Perrino, G. et al.. Feedback control promotes synchronisation of the cell-cycle across a population of yeast cells. In 2019 IEEE 58th Conference on Decision and Control (CDC) 933–938 (2019).

33. Garmendia-Torres, C., Tassy, O., Matifas, A., Molina, N. & Charvin, G. Multiple inputs ensure yeast cell size homeostasis during cell cycle progression. Elife 7, e34025 (2018).

34. Madden, K. & Snyder, M. Cell polarity and morphogenesis in budding yeast. Annu. Rev. Microbiol. 52, 687–744 (1998).

35. Kuramoto, Y. Chemical oscillations, waves, and turbulence. (Springer-Verlag, 1984).

36. Franklin, G. F., Powell, J. D. & Workman, M. L. Digital control of dynamic systems. vol. 3 (Addison-wesley Reading, MA, 1998).

37. Camacho, E. F. & Alba, C. B. Model predictive control. (Springer Science & Business Media, 2013).

