## Supplemental Information for "Cyber-yeast: Automatic synchronisation of the cell cycle in budding yeast through closed-loop feedback control"

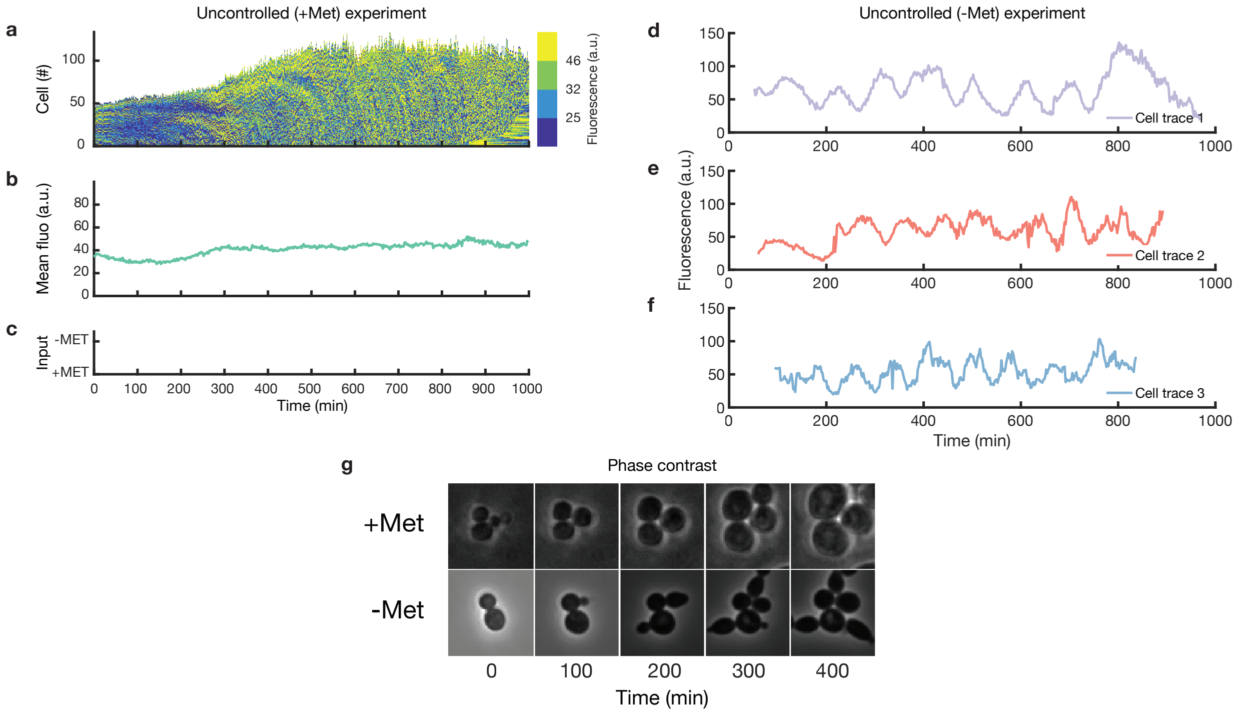


**Supplementary Figure 1. Experimental characterisation of the *non-cycling* strain. a-c**, Experimental characterisation of the non-cycling yeast strain grown in the automated microfluidics platform in methionine-rich medium (+MET). **a**, The number of cells and the distribution of YFP fluorescence intensity in the population over time. Fluorescence values are binned into 4 colours, corresponding to the quartiles, for clarity of visualisation. **b**, Average YFP fluorescence intensity in the cell population. **c**, Growth medium delivered to the cells as a function of time: +MET methionine-rich medium, -MET: methionine-depleted medium. **d-f**, YFP fluorescence intensity measured in three representative cells grown in methionine-depleted medium. **g**, Microscopy phase contrast images of cells grown in methionine-rich medium (top) or methionine-depleted medium (bottom) at the indicated time points.

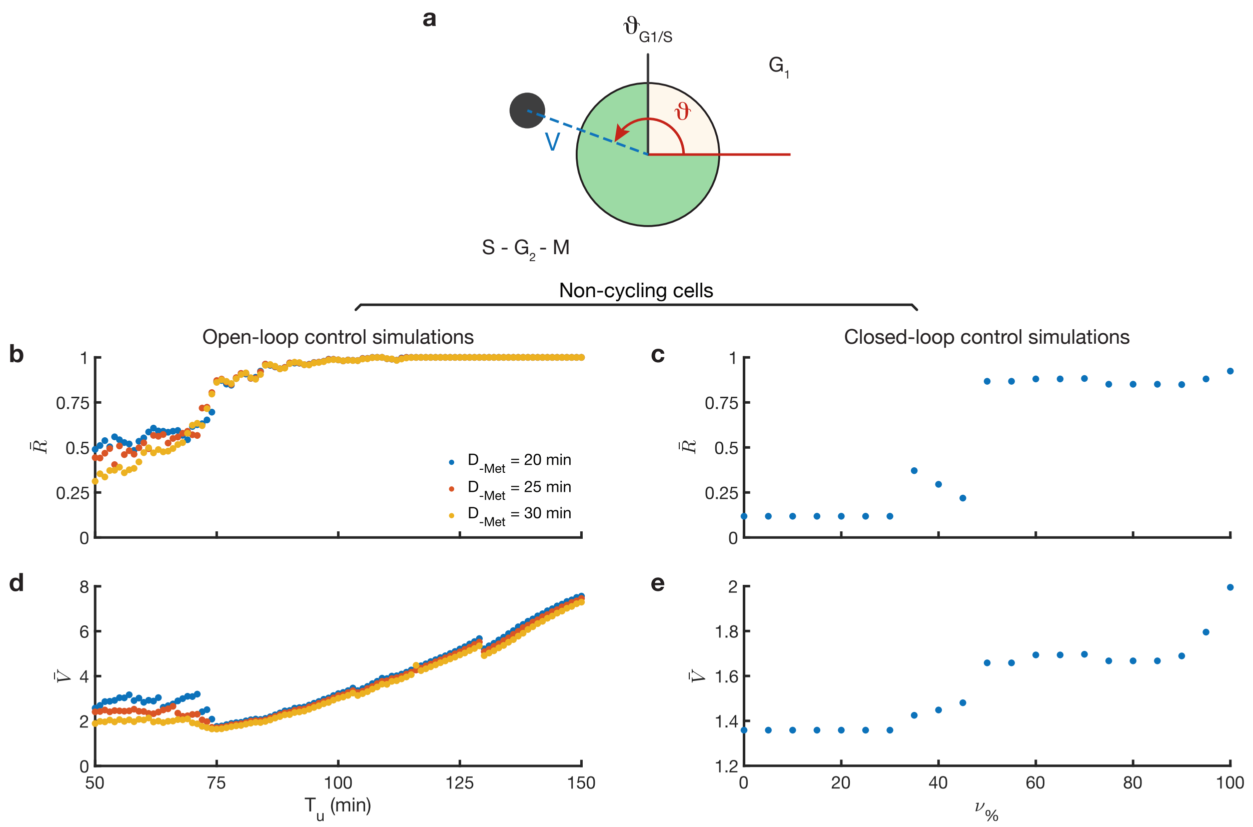


**Supplementary Figure 2. Mathematical modelling and simulation of the *non-cycling* strain. a**, Graphical representation of the mathematical model of the cell cycle. The black circle represents a cell evolving in $\mathbb{R}^2$. The cell cycle phase $\vartheta$ corresponds to the angle between the red line and the dashed blue line. The length of the dashed blue line corresponds to the cell volume. The yellow shaded sector corresponds to the $G_1$ phase, while the green shaded sector corresponds to the S–$G_2$–M phases. The cell cycle phase evolves counter clockwise. **b**, **d**, Average mean phase coherence $R$ and average volume growth $V$ for increasing values of the period $T_u$ and for three different pulse duration $D_{-Met}$ computed from numerical simulations of a population of cells in response to periodic stimulation with -MET pulses. **c**, **e**, The average mean phase coherence $R$ and average volume growth $V$ for increasing threshold values ($\nu_{\%}$) of the stop&go controller computed from numerical simulations of a population of non-cycling cells.

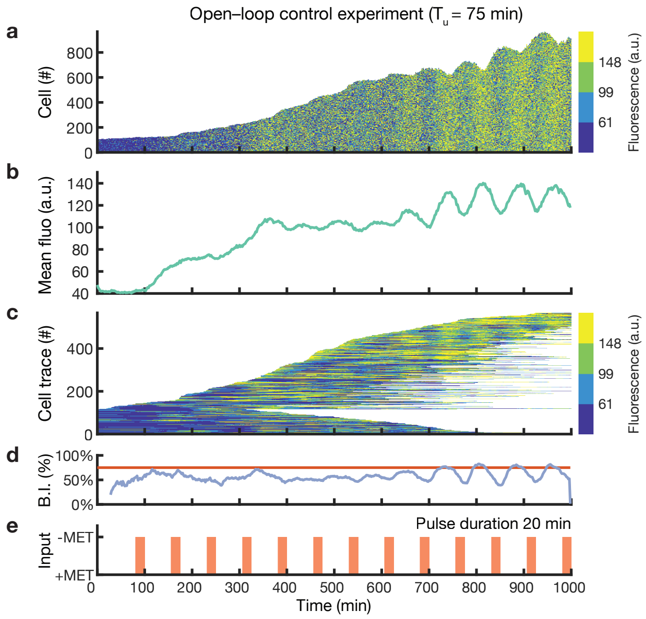


**Supplementary Figure 3. Additional *in vivo* open-loop control experiment in the *non-cycling* yeast strain with a shorter -MET pulse duration.** Non-cycling yeast strain cells grown in the automated microfluidics platform in response to alternating pulses of methionine-rich (+MET) and methionine-depleted (-MET) medium with period $T_u$ = 75 min and pulse duration $D_{-Met}$ = 20 min. **a**, The number of cells and the distribution of YFP fluorescence intensity in the population over time. Fluorescence values are binned into 4 colours, corresponding to the quartiles, for clarity of visualisation. **b**, Average YFP fluorescence intensity in the cell population. **c**, Single-cell fluorescence traces over time. Each horizontal line corresponds to one cell. Each line starts when the cell is first detected and ends when the cell exits the field of view. The number of tracked cells does not correspond to the total number of cells as only cells tracked for longer than 300 min are shown. **d**, Budding index (blue) reporting the percentage of cells in the budding phase (S-$G_2$-M) computed from the estimated cell cycle phases. The red line denotes the expected value of the budding index in the case of a totally desynchronised cell population. **e**, Growth medium delivered to the cells as a function of time: +MET methionine-rich medium, -MET: methionine-depleted medium.

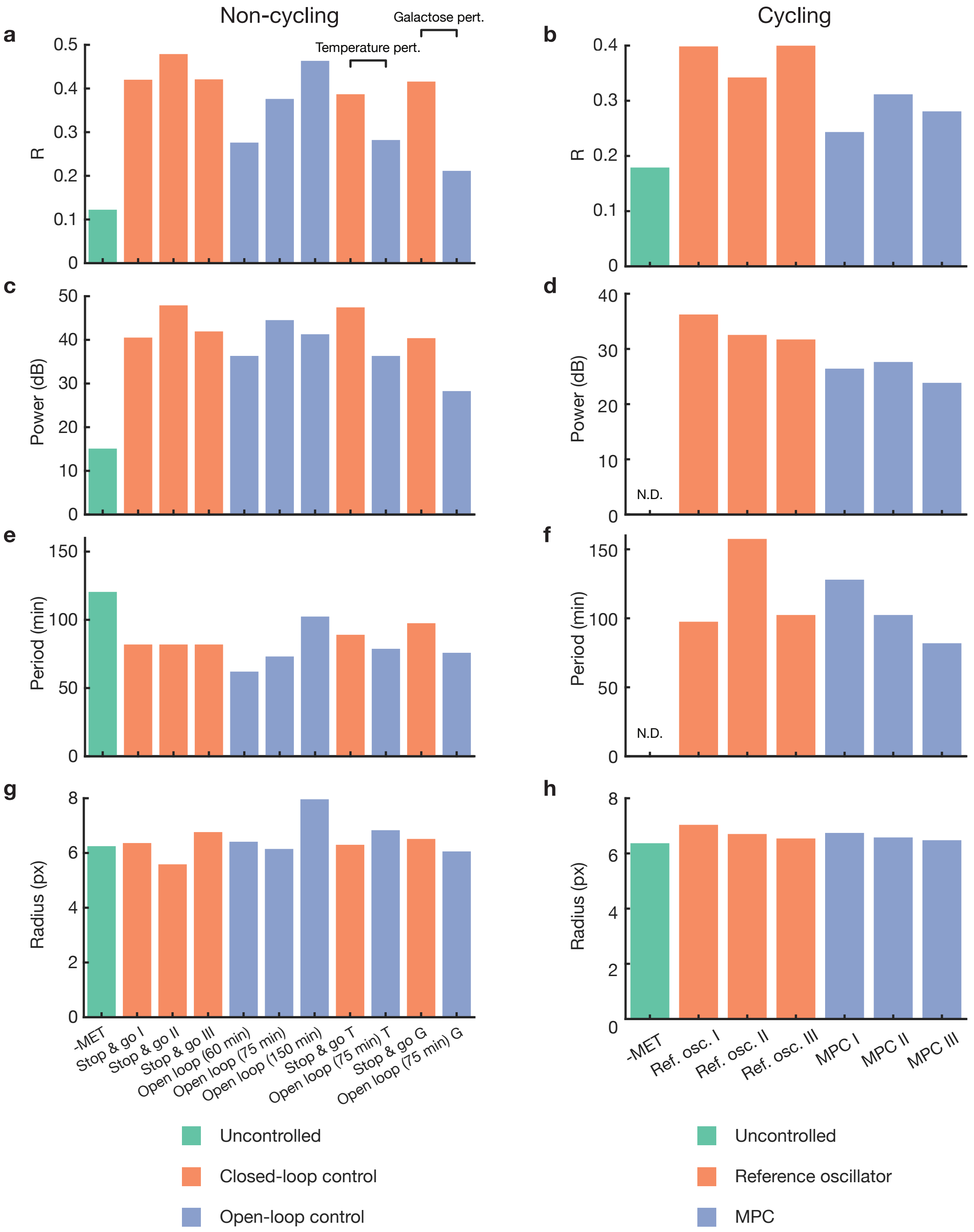


**Supplementary Figure 4. Quantification of cell cycle synchronisation and cell size. a**, **b**, Average phase coherence $R$ for all the experiments performed in the manuscript. The average was computed on the values between 330 and 530 min for the non-cycling strain, and between 100 and 600 min for the cycling strain. **c**, **d**, Amplitude of the leading peak of the power spectrum of the average YFP fluorescence intensity for the non-cycling (**c**) and the cycling (**d**) strains. **e**, **f**, Period of the power spectrum corresponding to the leading peak for the non-cycling (**c**) and the cycling (**d**) strains. **g**, **h**, Average radius of cells for the non-cycling (**g**) and the cycling (**h**) strain. *N.D.*: not determined – as the average YFP fluorescence intensity is not oscillatory and thus the estimation algorithm fails.

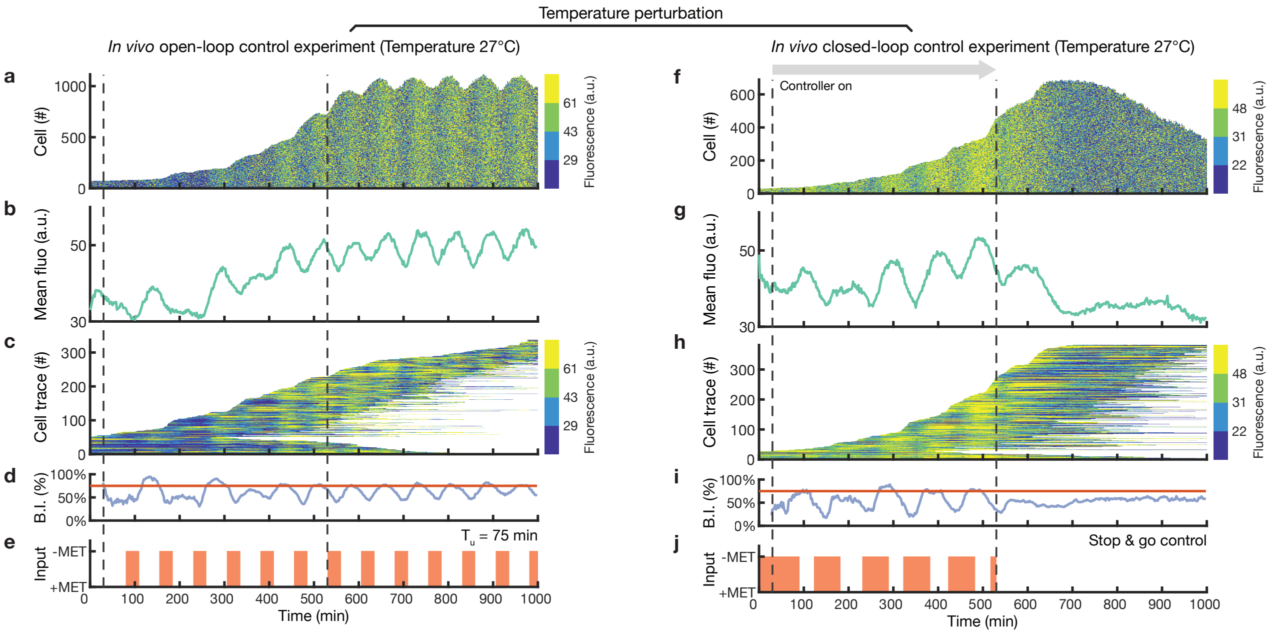


**Supplementary Figure 5. Closed-loop stop&go control of the *non-cycling* strain is robust to temperature perturbation.** Temperature was set to 27°C, rather than the nominal 30°C, to assess the robustness of the open-loop and closed-loop control strategies in perturbed environmental conditions. **a-e**, Open-loop control experiment: cells were stimulated with periodic -MET pulses of period $T_u$ = 75 min and pulse duration $D_{-Met}$ = 30 min. **f-j**, Closed-loop control experiment: an initial calibration phase of 30 min was required to set up the phase estimation algorithm. Dashed lines indicate the start and the end of the control experiment, after which cells are grown in methionine-rich medium. **a**, **f**, The number of cells and the distribution of YFP fluorescence intensity in the population over time. Fluorescence values are binned into 4 colours, corresponding to the quartiles, for clarity of visualisation. **b**, **g**, Average YFP fluorescence intensity in the cell population. **c**, **h**, Single-cell fluorescence traces over time. Each horizontal line corresponds to one cell. Each line starts when the cell is first detected and ends when the cell exits the field of view. The number of tracked cells does not correspond to the total number of cells as only cells tracked for longer than 300 min are shown. **d**, **i**, Budding index (blue) reporting the percentage of cells in the budding phase (S-$G_2$-M) computed from the estimated cell cycle phases. The red line denotes the expected value of the budding index in the case of a totally desynchronised cell population. **e**, **j**, Growth medium delivered to the cells as a function of time: +MET methionine-rich medium, -MET: methionine-depleted medium.

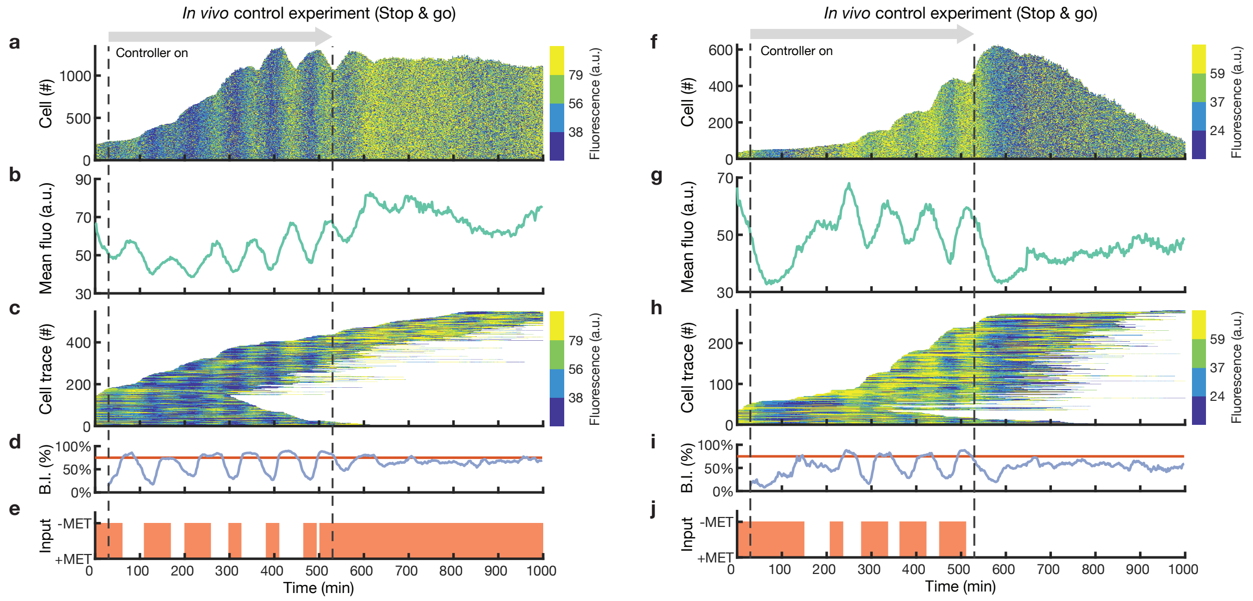


**Supplementary Figure 6. Closed-loop stop&go control experiments in the *non-cycling* yeast strain.** An initial calibration phase of 30 min was required to set up the phase estimation algorithm. Dashed lines indicate the start and the end of the control experiment, after which cells are grown either in methionine-depleted medium (**a**-**e**) or methionine-rich medium (**f**-**j**). **a**, **f** The number of cells and the distribution of YFP fluorescence intensity in the population over time. Fluorescence values are binned into 4 colours, corresponding to the quartiles, for clarity of visualisation. **b**, **g**, Average YFP fluorescence intensity in the cell population. **c**, **h**, Single-cell fluorescence traces over time. Each horizontal line corresponds to one cell. Each line starts when the cell is first detected and ends when the cell exits the field of view. The number of tracked cells does not correspond to the total number of cells as only cells tracked for longer than 300 min are shown. **d**, **i**, Budding index (blue) reporting the percentage of cells in the budding phase (S-$G_2$-M) computed from the estimated cell cycle phases. The red line denotes the expected value of the budding index in the case of a totally desynchronised cell population. **e**, **j**, Growth medium delivered to the cells as a function of time: +MET methionine-rich medium, -MET: methionine-depleted medium.

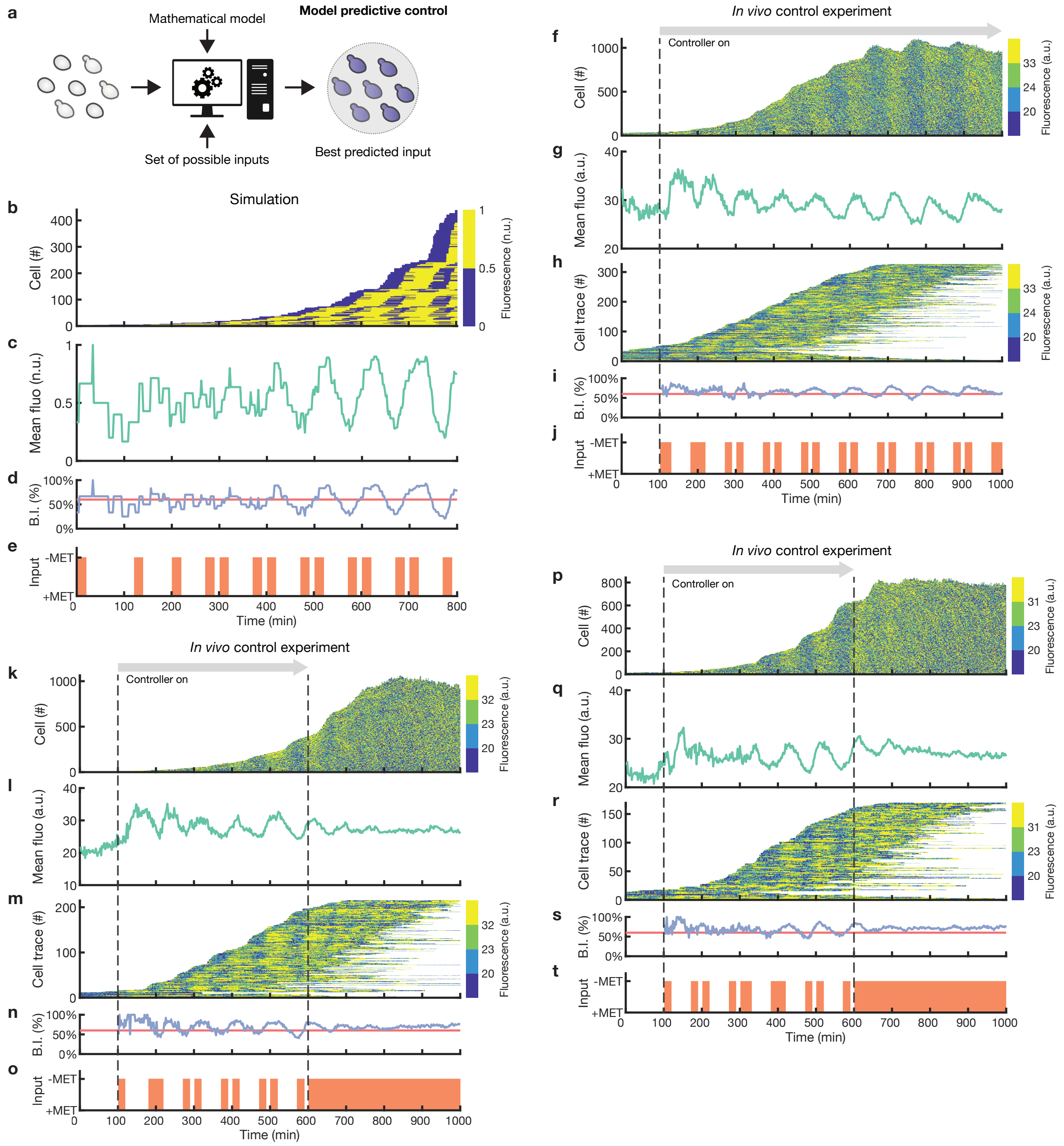


**Supplementary Figure 7. Closed-loop model predictive control (MPC) experiments in the *cycling* yeast strain. a**, Schematic illustration of the Model predictive control (MPC) strategy. The controller uses a mathematical model to predict the future behaviour of the cell cycle across the population of cells to find the best sequence of -MET pulses to synchronise the cells. **b-e**, Numerical simulation of the MPC control strategy. **f-t**, Experimental implementation of the MPC strategy. An initial calibration phase of 100 min is required to set up the phase estimation algorithm. Dashed lines indicate the start and the end of the control experiment, after which cells are grown in methionine-depleted medium. **b**, **f**, **k**, **p**, The number of cells and the distribution of YFP fluorescence intensity in the population over time. Fluorescence values are binned into 4 colours, corresponding to the quartiles, for clarity of visualisation. **c**, **g**, **l**, **q**, Average YFP fluorescence intensity in the cell population. **h**, **m**, **r**, Single-cell fluorescence traces over time. Each horizontal line corresponds to one cell. Each line starts when the cell is first detected and ends when the cell exits the field of view. The number of tracked cells does not correspond to the total number of cells as only cells tracked for longer than 300 min are shown. **d**, **i**, **n**, **s**, Budding index (blue) reporting the percentage of cells in the budding phase (S-$G_2$-M) computed from estimated cell cycle phases. Red line denotes the expected value of the budding index in the case of a totally desynchronised cell population. **e**, **j**, **o**, **t**, Growth medium delivered to the cells as a function of time: +MET methionine-rich medium, -MET: methionine-depleted medium.

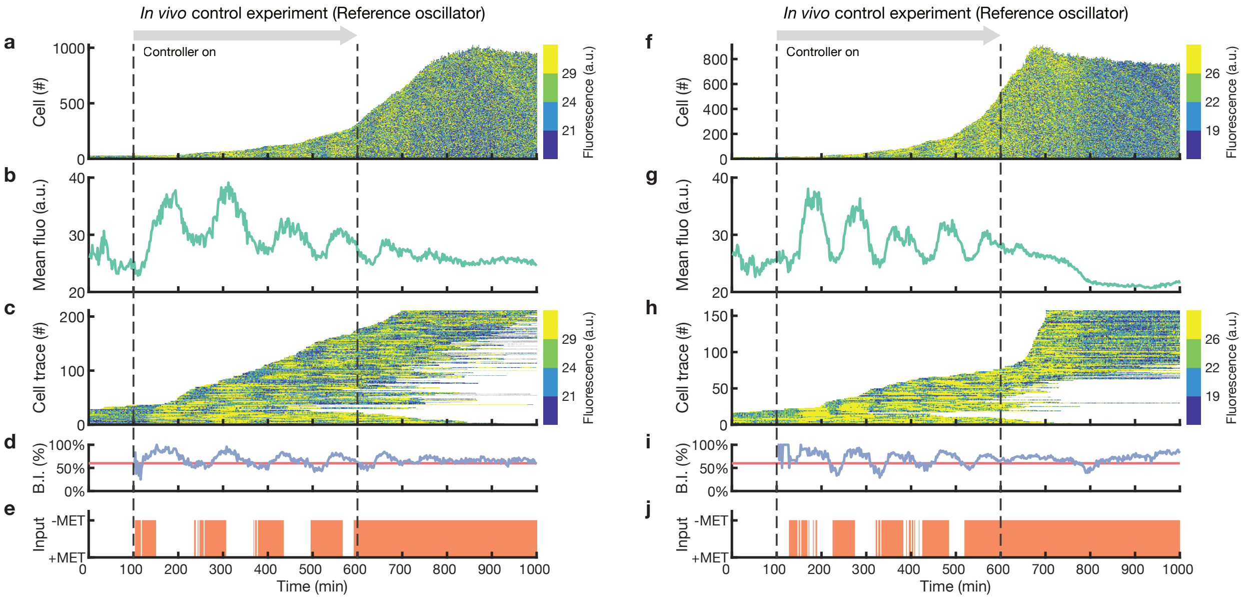


**Supplementary Figure 8. Closed-loop reference oscillator control experiments in the *cycling* yeast strain.** Experimental implementation of the reference oscillator control strategy. An initial calibration phase of 100 min is required to set up the phase estimation algorithm. Dashed lines indicate the start and the end of the control experiment, after which cells are grown in methionine-depleted medium. **a**, **f**, The number of cells and the distribution of YFP fluorescence intensity in the population over time. Fluorescence values are binned into 4 colours, corresponding to the quartiles, for clarity of visualisation. **b**, **g**, Average YFP fluorescence intensity in the cell population. **c**, **h**, Single-cell fluorescence traces over time. Each horizontal line corresponds to one cell. Each line starts when the cell is first detected and ends when the cell exits the field of view. The number of tracked cells does not correspond to the total number of cells as only cells tracked for longer than 300 min are shown. **d**, **i**, Budding index (blue) reporting the percentage of cells in the budding phase (S-$G_2$-M) computed from estimated cell cycle phases. Red line denotes the expected value of the budding index in the case of a totally desynchronised cell population. **e**, **j**, Growth medium delivered to the cells as a function of time: +MET methionine-rich medium, -MET: methionine-depleted medium.

**Supplementary Table 1. Yeast strains used in this study. All strains are W303 congenic.**

| Strain | Description | Genotype | Source |
|---|---|---|---|
| SJR14a4d | Non-cycling strain | cln1Δ, cln2Δ::CLN2p-Venus::TRP1, cln3Δ::LEU2, trp1Δ::TRP1::MET3p-CLN2, HTB2-mCherry::HIS5 | Ref.[1] |
| GC84-35B | Background of cycling strain | cln3Δ::LEU2, CDC10-YFP::LEU2, WHI5-GFP::KanMx, TRP1::MET3p-CLN2 | Ref.[2] |
| yDdB028 | Cycling strain | cln3Δ::LEU2, CDC10-YFP::LEU2, WHI5-GFP::KanMx, TRP1::MET3p-CLN2, HTB2-mCherry::natNT2 | This study |

**Supplementary Table 2. Plasmids used in this study.**

| Plasmid | Type | Source |
|---|---|---|
| pRS41N | Replicative | Euroscarf; ref.[3] |
| pRS41N-GAP-CYC | Replicative | C. Wilson |
| pRS41N-GAP-HTB2-mCherry-CYC | Replicative | This study |

### SUPPLEMENTARY NOTES

**Supplementary Note 1. Derivation of the reference oscillator strategy.** The reference oscillator strategy was derived from the one proposed by Bai and Wen to asymptotically synchronise a homogeneous population of phase oscillators using a single (i.e., common) control input[4]. To derive the reference oscillator strategy, we made a series of assumptions that allowed us to adapt the state feedback control law proposed by Bai and Wen[4] to solve the cell cycle synchronisation problem in a population of cycling yeast cells using an asymmetrically bounded input. Firstly, we neglected the dynamics associated to cell growth and cell division. Such an assumption implies a constant number of cycling cells in the population. Secondly, we assumed that the cell cycle dynamics in each cell is well approximated by a first-order reduced phase dynamics determined by a phase response curve (PRC). Such an assumption is well in line with our mathematical framework (see Modelling section in Methods). Let $N$ denote the number of cycling cells in the cell population. Then, the cell cycle dynamics of each cell is represented by an ODE model that describes the cell cycle phase evolution in time:

$$\frac{d}{dt}\vartheta = \omega + z(\vartheta)\, u\,,$$

where $\omega \in \mathbb{R}_+$ is the natural frequency of the cell cycle oscillator, $\vartheta \in \mathbb{S}^1$ is the 2π-periodic cell cycle phase on the unit circle, $u \in \{u_{OFF}, u_{ON}\}$ is the external trigger input, and $z : \mathbb{S}^1 \to \mathbb{R}_+$ is the 2π-periodic PRC function that models the linear response of the cell cycle phase $\vartheta$ to the input $u$. Note that the proposed control input $u$ is asymmetrically bounded *a priori*. Indeed, in our modelling framework $u_{OFF} = 0$ (i.e., when the cell is fed with methionine-rich medium), while $u_{ON} = 1$ (i.e., when the cell is fed with methionine-depleted medium). Also, in our modelling framework, we mathematically described the PRC function as:

$$z(\vartheta) = \begin{cases} \omega_z & if\ 0 \le \vartheta < \vartheta_{G_1/S} \\ 0 & if\ \vartheta_{G_1/S} \le \vartheta < 2\pi \end{cases},$$

where $\omega_z \in \mathbb{R}_+$ is the angular velocity added to the cell cycle phase dynamics when $u = 1$ (i.e., $u = u_{ON}$), and $\vartheta_{G_1/S}$ is the cell cycle phase value at the $G_1$ to S transition. We then derived a state feedback control law for an asymmetrically bounded input as presented in the Section III of Bai and Wen[4]. We introduced a *reference* oscillator modelled by the first-order reduced phase dynamics:

$$\frac{d}{dt}\vartheta_r = \omega_r + \gamma \sum_{m=1}^{N} \sin\left(\vartheta_m - \vartheta_r\right),$$

where $\omega_r \in \mathbb{R}$ is the natural frequency of the reference oscillator, $\vartheta_r \in \mathbb{S}^1$ is the 2π-periodic cell cycle phase of the reference oscillator, $\gamma \in \mathbb{R}_+$ is the coupling strength, and $\vartheta_m$ is the cell cycle phase of the *m*-th cell in the population. Let $\vartheta = [\vartheta_1, \ldots, \vartheta_N]^T$ and $h(\vartheta) = [z(\vartheta_1), \ldots, z(\vartheta_N)]^T$. Let $\delta_m = \vartheta_m - \vartheta_r$ define the phase error associated to the *m*-th cell. Let $\delta = [\delta_1, \ldots, \delta_N]^T$ and $sin(\delta) = [sin(\delta_1), \ldots, sin(\delta_n)]^T$. We thus considered the state feedback control law [refer to Section III in Bai and Wen[4]]:

$$u = -u_{ON}\, \rho_u\, (sin(\delta)^T\, h(\vartheta))\,,$$

where the continuous function $\rho_u(\cdot) : \mathbb{R} \to [-1,0]$ satisfies $\rho_u(s) = 0,\ \forall s \geq 0$ and $-s \geq \rho_u(s)\, s > 0$, $\forall s < 0$. Being that $\|sin(\delta)^T\, h(\vartheta)\| \leq N\, \omega_z$, then we could choose:

$$\rho_u(s) = \begin{cases} 0 & if\ s \geq 0 \\ -\dfrac{s}{N\, \omega_Z} & if\ s < 0 \end{cases},$$

where $s = sin(\delta)^T\, h(\vartheta)$. However, $u \in \{0,1\}$ is a bang-bang control input, thus we devised a robust realisation of the continuous function $\rho_u(\cdot)$, that is:

$$\rho_u(s) = \begin{cases} 0 & if\ s \geq 0 \\ -1 & if\ s < 0 \end{cases},$$

where $s = sin(\delta)^T\, h(\vartheta)$. The previous function leads to the bang-bang state feedback control:

$$u = \begin{cases} 0 & if\ s \geq 0 \\ 1 & if\ s < 0 \end{cases},$$

where $s = sin(\delta)^T\, h(\vartheta)$. However,

$$sin(\delta)^T\, h(\vartheta) = \sum_{m=1}^{N} \sin(\vartheta_m - \vartheta_r)\; z(\vartheta_m) = \sum_{m=1}^{N} \mathrm{a_m} \sin(\vartheta_m - \vartheta_r)\; \omega_z = \omega_z \sum_{m=1}^{N} \mathrm{a_m} \sin(\vartheta_m - \vartheta_r)\,,$$

where $\mathrm{a_m} \in \{0,1\}$ is a coefficient denoting if the *m*-th cell is in the $G_1$ phase, that is:

$$a_m = \begin{cases} 1, & if\ 0 \leq \vartheta_m < \vartheta_{G_1/S} \\ 0, & if\ \vartheta_{G_1/S} \leq \vartheta_m < 2\pi \end{cases}.$$

Thus, we obtained that

$$s = sin(\delta)^T\ h(\vartheta) \geq 0\ \Leftrightarrow \omega_z \sum_{m=1}^{N} \mathrm{a_m} \sin(\vartheta_m - \vartheta_r) \geq 0\ .$$

Being $\omega_z > 0$, then the previous inequality holds also when

$$\sum_{m=1}^{N} \mathrm{a_m} \sin(\vartheta_m - \vartheta_r) \geq 0\ .$$

Finally, we obtained the bang-bang state feedback control law as reported in the Methods:

$$u = \begin{cases} u_{OFF} = 0, & if \sum_{m=1}^{N} \mathrm{a_m} \sin(\vartheta_m - \vartheta_r) \geq 0 \\ u_{ON} = 1, & if \sum_{m=1}^{N} \mathrm{a_m} \sin(\vartheta_m - \vartheta_r) < 0 \end{cases}.$$

Such a control law, under the previous assumptions, guarantees that the cell cycle phases converge asymptotically towards the reference phase according the Theorem 2 stated in Bai and Wen[4].

### SUPPLEMENTARY MOVIES

**Supplementary Movie 1.** Experimental characterisation of the non-cycling strain in methionine-depleted medium. Related to Fig. 2a-e.

**Supplementary Movie 2.** Open-loop control of non-cycling cells with forcing period $T_u$ = 75 min and pulse duration $D_{-Met}$ = 30 min. Related to Fig. 2k-o.

**Supplementary Movie 3.** Open-loop control of non-cycling cells with forcing period $T_u$ = 150 min and pulse duration $D_{-Met}$ = 30 min. Related to Fig. 2p-t.

**Supplementary Movie 4.** Closed-loop control of non-cycling cells. Related to Fig. 3f-j.

**Supplementary Movie 5.** Experimental characterisation of the cycling strain in methionine-supplemented medium. Related to Fig. 4f-j.

**Supplementary Movie 6.** Closed-loop control of cycling cells using the refence oscillator strategy. Related to Fig. 4p-t.

### SUPPLEMENTARY REFERENCES


1. Rahi, S. J., Pecani, K., Ondracka, A., Oikonomou, C. & Cross, F. R. The CDK-APC/C Oscillator Predominantly Entrains Periodic Cell-Cycle Transcription. *Cell* **165**, 475–487 (2016).
2. Charvin, G., Cross, F. R. & Siggia, E. D. Forced periodic expression of G1 cyclins phase-locks the budding yeast cell cycle. *Proc. Natl. Acad. Sci. U. S. A.* **106**, 6632–6637 (2009).
3. Taxis, C. & Knop, M. System of centromeric, episomal, and integrative vectors based on drug resistance markers for Saccharomyces cerevisiae. *Biotechniques* **40**, 73–78 (2006).
4. Bai, H. & Wen, J. T. Asymptotic Synchronization of Phase Oscillators With a Single Input. *IEEE Trans. Automat. Contr.* **64**, 1611–1618 (2019).
