## Supplementary Data 1 for "Cyber-yeast: Automatic synchronisation of the cell cycle in budding yeast through closed-loop feedback control": code_Supplementary_Data_1.m

```
%% code_Supplementary_Data_1.m
%%% OCTOBER 21, 2020

%% code_Supplementary_Data_1.m estimates the cell cycle period of the cycling strain

clear all

close all


%% Load the single-cell traces
load('./Supplementary_Data_1.mat')


%% Set the sampling time
sampling_time = 3;


%% Set the total number of frames and cell traces
[dim_exp, dim_traces] = size(cell_traces);


%% Allocate matrices and vectors
norm_traces = NaN(dim_exp, dim_traces);
square_traces = NaN(dim_exp, dim_traces);

vct_D_g1 = [];
vct_D_sg2m = [];


%%
for z = 1:dim_traces
    
    %% Normalisation
    norm_traces(:,z) = (cell_traces(:,z) - min(cell_traces(:,z))) / ...
        (max(cell_traces(:,z)) - min(cell_traces(:,z)));
    
    
    %% Binarisation
    square_traces(:,z) = norm_traces(:,z) >= 0.3;
    
    
    %% Improving square trace - First step
    wrong = find((square_traces(3:end-2,z) ~= square_traces(1:end-4,z) & ...
        square_traces(3:end-2,z) ~= square_traces(2:end-3,z) & ...
        square_traces(3:end-2,z) ~= square_traces(4:end-1,z) & ...
        square_traces(3:end-2,z) ~= square_traces(5:end,z)));
    
    square_traces(2+wrong,z) = (~square_traces(2+wrong,z));
    
    for v=1:2
        
        wrong = find((1*(square_traces(3:end-2,z) ~= ...
            square_traces(1:end-4,z)) + (square_traces(3:end-2,z) ~= ...
            square_traces(2:end-3,z)) + (square_traces(3:end-2,z) ~= ...
            square_traces(4:end-1,z)) + (square_traces(3:end-2,z) ~= ...
            square_traces(5:end,z))) >= 3);
        
        square_traces(2+wrong,z) = (~square_traces(2+wrong,z));
        
    end
    
    
    %% Improving square trace - Second step
    a = diff(square_traces(:,z));
    
    b = find(a);
    
    if length(b) >= 2
        
        c = find(diff(b) <= 8 & a(b(2:end)) == -1);
        
        for q=1:length(c)
            
            square_traces(b(c(q))+1:b(c(q)+1),z) = square_traces(b(c(q)),z);
        
        end
        
    end
    
    
    %% Compute durations of G1 and S-G2-M phases for z-th cell
    a = diff(square_traces(:,z));
    b = find(a);
    c = diff(b);
    
    
    tmp_g1 = sampling_time * c(a(b(1:end-1))==-1);
    tmp_sg2m = sampling_time * c(a(b(1:end-1))==1);
    
    
    vct_D_g1 = [vct_D_g1; tmp_g1];
    vct_D_sg2m = [vct_D_sg2m; tmp_sg2m];
    
end

%% Durations of G1 and S-G2-M phases
D_g1 = median(vct_D_g1);
D_sg2m = median(vct_D_sg2m);

T = D_g1 + D_sg2m;


%% Recap table
VarNames = {'T','T_g1','T_sg2m'};

Tab = table(T, D_g1, D_sg2m, 'RowNames', {'+MET'}, 'VariableNames', VarNames);

disp(Tab);

%% Boxplot
F = figure('Position', [1 1 480 480], 'DefaultAxesFontSize', 16);

data_var = [vct_D_g1;vct_D_sg2m];

group_var = [repmat({'G1 phase'}, length(vct_D_g1), 1); ...
    repmat({'S-G2-M phases'}, length(vct_D_sg2m), 1)];

boxplot(data_var, group_var);

ylim([0,150]);
ylabel('Duration (min)')

% Print the boxplot
print(F, './Supplementary_Data_1', '-dpng');
```
