## Supplementary figures and images for "Cyber-yeast: Automatic synchronisation of the cell cycle in budding yeast through closed-loop feedback control"

### Supplementary_Data_1.png

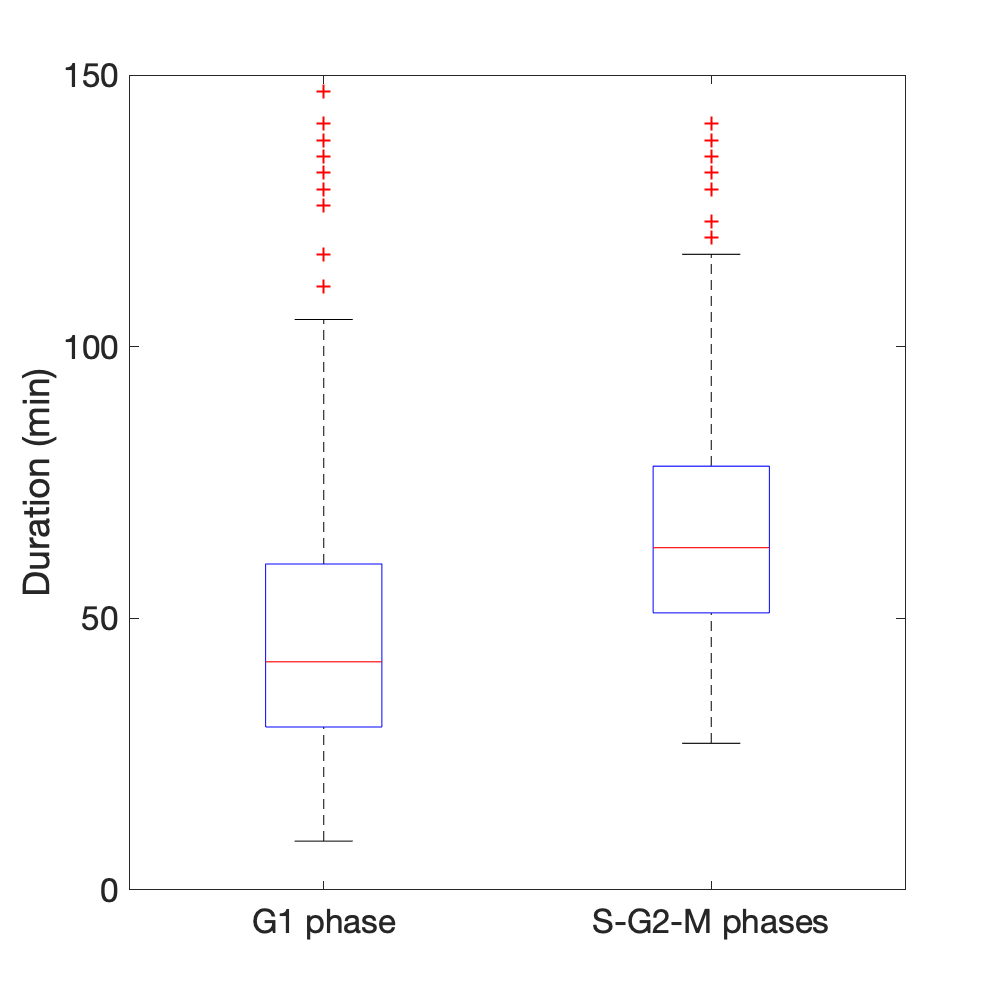
